# A Genome-Scale Metabolic Model of Marine Heterotroph *Vibrio splendidus* sp. 1A01

**DOI:** 10.1101/2022.04.15.488298

**Authors:** Arion Iffland-Stettner, Hiroyuki Okano, Matti Gralka, Ghita Guessous, Kapil Amarnath, Otto X. Cordero, Terence Hwa, Sebastian Bonhoeffer

## Abstract

While the *Vibrio splendidus* species is best known as an opportunistic pathogen in oysters, the *Vibrio splendidus* sp. 1A01 strain was first identified as an early colonizer of synthetic chitin particles incubated in seawater. To gain a better understanding of its metabolism, a genome-scale metabolic model (GSMM) of *V. splendidus* sp. 1A01 was reconstructed. GSMMs enable us to simulate all metabolic reactions in a bacterial cell using Flux Balance Analysis. A draft model was built using an automated pipeline from BioCyc. Manual curation was then performed based on experimental data, in part by gap-filling metabolic pathways and tailoring the model’s biomass reaction to *V. splendidus* sp. 1A01. The challenges of building a metabolic model for a marine microorganism like *V. splendidus* sp. 1A01 are described.

## Introduction

The heterotrophic, Gram-negative *Vibrio splendidus* species is found ubiquitously in the ocean, both in close association with marine animals (like bivalves^1–3^) and as an “environmental” microorganism in marine microbial communities^4,5^. When associated with marine animals, its interactions with the host can be either commensal^6^ or pathogenic^7^. As a pathogen, *V. splendidus* induces vibriosis^8^ and is of relevance to the aquaculture industry, causing outbreaks in hatcheries around the world, at great economic cost^2^. *V. splendidus* infects, for example, farmed oysters like *C. gigas*^2^ and *C. virginica*^3^, as well as farmed sea cucumbers like *A. japonicus*^9–11^. As an “environmental” microorganism, *V. splendidus* is found both in seawater and coastal sediments^12^, its lifestyle ranging from planktonic^13^ to particle-associated^5^. It is as a particle-associated marine microorganism, with an ecological role to play in establishing microbial communities on ocean particles, that the *V. splendidus* sp. 1A01 strain was first isolated^5^. More precisely, *V. splendidus* sp. 1A01 was isolated from synthetic chitin particles incubated in seawater samples to allow for the assembly of marine microbial communities^5^. *V. splendidus* sp. 1A01 was identified as an early colonizer of the chitin particles, secreting enzymes that break down chitin and, thereby, lay the groundwork for microbial community assembly^5^. More recently, *V. splendidus* sp. 1A01 was used to study how the metabolism of specific heterotrophic marine bacteria affects organic matter remineralization^14^.

Genome-scale metabolic models (GSMMs) have proven to be powerful tools in systems biology for simulating the metabolism of bacteria such as *E. coli*^15–17^. Mathematically, a GSMM is comprised of a stoichiometric matrix encoding all of the reactions in a cell, in addition to exchange fluxes with the environment^15,18^. The model includes also a biomass reaction that converts, in experimentally measured proportions, the building blocks of a cell (*e*.*g*. nucleic acids, amino acids, vitamins, cofactors) into biomass^15,18^. Finally, upper and lower bounds are imposed on the permissible flux through every reaction^15,18^. Thus, the model captures the full metabolic capabilities of a microorganism for growth conditions where measurements are available or can be extrapolated.

Constraint-based computational methods like Flux Balance Analysis (FBA) can be used to simulate the distribution of fluxes through a whole-cell metabolic network, when supplied with an objective function and constraints^15,18^. Often, the objective function being optimized under steady-state exponential growth is biomass production^15,18^, while a constraint can be the measured carbon uptake rate^15,18^. If so, FBA will calculate the optimal flux through the cell’s biomass reaction given this constraint. By virtue of its implementation through linear programming, FBA is computationally inexpensive^18^. Beyond calculating optimal growth rates, FBA has been successfully applied in the context of metabolic engineering^19–21^, identifying drug targets^22–24^, studying the properties of metabolic networks^25^, simulating bacterial growth in three dimensions^26^, and predicting cross-feeding interactions within microbial communities^27–29^. Finally, metabolic models can be integrated with a variety of omics data, including metabolomics^30–32^, transcriptomics^33–35^, and proteomics^34,36^.

To facilitate the study of *V. splendidus* sp. 1A01, a GSMM was reconstructed, the first for a *Vibrio splendidus* strain (Fig. 1). Reconstruction began by feeding the annotated genome of *V. splendidus* sp. 1A01 into an automated pipeline from BioCyc (a database of metabolic reactions^37^) to obtain a draft metabolic model. Limited by the well-known incompleteness of genomic annotation (recent estimates of the average annotation completeness for bacterial genomes range from 52% to 79%, depending on the annotation method^38^), the draft metabolic model had to undergo extensive manual curation. First, based on the growth of *V. splendidus* sp. 1A01 on a wide variety of carbon sources, metabolic pathways were gap-filled^39^. Second, the proportions in which cellular building blocks are converted into *V. splendidus* sp. 1A01 biomass were experimentally measured, to curate the model’s biomass reaction. Third, the GAM (Growth-Associated Maintenance^40^) and NGAM (Non-Growth-Associated Maintenance^40^) of *V. splendidus* sp. 1A01 were also measured. The curated model was then quantitatively validated, using still more experimental data. Along the way, building a model for a marine microbe like *V. splendidus* sp. 1A01 presented a number of technical challenges, which are addressed in the Conclusion.

**Figure 1.**
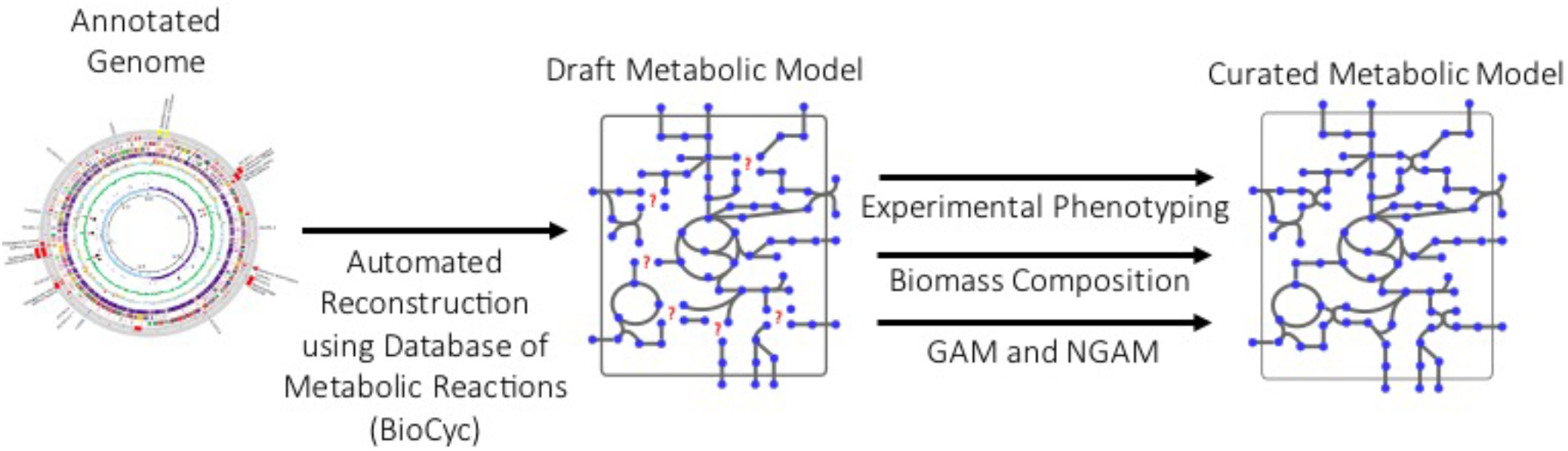
Computational Pipeline for Building the Metabolic Model of *V. splendidus* sp. 1A01.

## Methods

### Culture conditions for quantitative measurements

The growth media used were marine broth (Difco Marine Broth 2216) and a complete minimal medium for growing copiotrophic, heterotrophic marine bacteria. Marine broth was prepared by dissolving 37.4 g/L in ddH_2_O, boiling for 1 min, and filtering through a 0.22-μm filter for sterilization. It was stored at room temperature. The minimal medium consisted of a carbon source, 10 mM NH_4_Cl, 0.5 mM Na_2_HPO_4_, 1 mM Na_2_SO_4_, simple salts to mimic seawater (0.343 M NaCl, 14.75 mM MgCl_2_ · 6H_2_O, 4 mM CaCl_2_ · 2H_2_O, and 27 mM KCl, which is designated as 1x SW), 40 mM HEPES pH 8 as the buffer, and trace metals such as iron. All components were filter-sterilized using a 0.22-μm filter. The minimal medium was stored at 4°C. For a full description of the preparation and composition of the minimal medium, see ref. 41.

Preparing batch cultures used for measuring RNA, protein, metabolites, and cell dry weight of 1A01 involved three steps: 1) a seed culture, 2) a preculture, and 3) a batch (experimental) culture. The seed culture was started by inoculating 2 mL of marine broth in a 16 mm × 125 mm test tube (borosilicate glass, Fisherbrand, Cat. No. 14-961-30) from a single colony on a marine broth/agar plate. Once the seed culture saturated (which took ∼7 hrs), the cells were washed and resuspended in 1x SW to an OD_600_ of ∼1 before being diluted into the experimental medium (3 mL in a 18 mm × 150 mm tube) for growth overnight, such that, by the following day, the preculture a) doubled ≥10 times and b) remained growing exponentially. While the preculture was still in exponential growth, we diluted the preculture into fresh experimental medium prewarmed to 27°C to an OD_600_ of 0.01 - 0.02. All cultures (seed, pre-, and batch) were grown in a water bath shaker at 27°C with shaking at 250 rpm. OD_600_ was measured using a Thermo Scientific GENESYS 20 or 30 Spectrophotometer.

### Culture conditions for growth phenotype screening

High-throughput screening was also performed to obtain a coarse growth phenotype for *V. splendidus* sp. 1A01. After thawing (at room temperature) a frozen culture sample (5% DMSO), 20 μL of the stock culture was transferred to 180 μL marine broth in 96-well plates and grown for 94 hours at room temperature without shaking. The cultures were diluted 1:1 in carbon-free minimal MBL medium (see Doc. S1 for full recipe) for 2 hours, then transferred into 384-well plates filled with 70 μL minimal medium per well and one of 78 added carbon sources (see Table S1 for full list) using a pinning tool (V&P Scientific VP 408, 0.2 μL hanging drop volume). The concentration of carbon atoms was normalized to 40 mM for each carbon source. All plates were then covered with transparent film (Life Technologies MicroAmp Optical Adhesive Film) and stored in the dark at room temperature. All plates were read at least once a day for ten days using a stack plate reader (Tecan Spark) to measure optical density at 600 nm.

### RNA, protein, and amino acid measurements

RNA and proteins were measured as described before^42^, with modifications. Since the cells from cultures grown on glucose, GlcNAc, and glucosamine were not harvested well by the short centrifugation employed in the regular protocol, they were first chilled on ice-water for 5 minutes, and harvested by centrifugation at 15,000 rpm (Eppendorf Centrifuge 5424) for 15-20 min at 4°C. For protein measurements, the cells were further rinsed in the minimal medium lacking carbon, nitrogen, and phosphorus sources and harvested by centrifugation for 15-20 min. The effects of the long centrifugation at 4°C were assessed by comparing measured protein concentrations with those using the regular short centrifugation for cells grown on glycerol, since glycerol cultures were harvested well by the regular short centrifugation. Little difference (<2%) was observed.

For RNA measurements, 1.5 mL of an exponentially growing culture was pelleted, fast-frozen on dry ice and stored. Pellets were thawed and washed twice with 0.7 M cold HClO_4_ then digested for 60 minutes at 37°C using 300 μL of 0.3 M KOH. Samples were periodically stirred. The cell extracts were then neutralized with 100 μL of 3 M HClO_4_ and centrifuged at 13,000 rpm for 3 minutes. The soluble fraction was collected and the remaining pellets washed twice with 550 μL of 0.5 M HClO_4_. The resulting final volume of 1.5 mL was centrifuged once more to eliminate remaining debris and its absorbance at 260 nm was measured using a Bio-Rad spectrophotometer. The RNA concentration was determined as OD_260_*31/OD_600_, where the conversion factor is based on RNA’s extinction coefficient.

Protein amounts were quantified using the Biuret method. 1.5 mL of an exponentially growing culture was pelleted, washed with 1x SW, resuspended in 200 μL of 1x SW and fast frozen on dry-ice. The cell pellet was then thawed at room temperature. 100 μL of 3 M NaOH was added to the pellet and samples were incubated on a heat block at 100°C for 5 minutes to hydrolyze the proteins. Protein amounts in the samples were determined using the Biuret method. 100 μL of 1.6% CuSO_4_ was added to the protein extracts and samples were centrifuged for 3 minutes at 13,000 rpm. The absorbance of the soluble fraction was read at OD_555_ using a spectrophotometer. A series of 200 μL BSA standards were taken through the same procedure to get a standard curve.

Glutamate and glutamine pools were measured with the no-harvest protocol as described previously^43,44^.

### Cell dry weight measurements

In a regular protocol to measure cell dry weight (CDW), a cell pellet is washed with water to remove salts from the culture medium. However, such a protocol cannot be applied to bacterial cells grown at high osmolarity because the cells are lysed in water. Therefore, we had to develop a novel protocol to estimate the CDW of bacterial cells grown at high osmolarity.

A *V. splendidus* sp. 1A01 culture was grown to an OD_600_ of 0.5, chilled on ice-cold water for 5-15 minutes, and harvested by centrifugation at 4°C. The cell pellet was washed with the same volume of cold NaCl solution as the culture, and washed again with 40 mL of the same NaCl solution. Concentrations of NaCl were 0.45 M for cultures grown at 1x SW and 0.52 M for cultures grown at 1.25x SW or in marine broth. The density of the washing NaCl solution was measured with water as a reference. After the wet pellet was weighed, it was suspended in water, transferred to a weighing dish, and dried in an oven at 85-95°C until the weight became stable, typically for three days. The dry pellet was weighed within 10 seconds before it absorbed water and thereby significantly increased in weight. CDW was estimated as follows. Let *x* and *y* be the weights of wet and dry cell pellets, respectively, *ρ*the weight of NaCl per the weight of water in the washing solution, *w* the harvested amount of cells in units of OD_600_∙mL, and *z* the weight of extracellular water in the wet pellet.

With these parameters, we can represent *α*, the CDW per OD_600_∙mL, as a function of *β*, the cellular water weight per CDW.

From the equality of CDW,

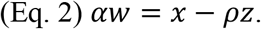

From the equality of water weight present in the wet pellet,

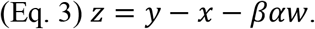

From the above two equations, we obtain

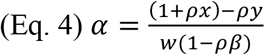

In *E. coli, β* is reported to be *∼*2 mg/mg CDW^45^. Like *E. coli, V. splendidus* sp. 1A01 is a rod-shaped, gram-negative bacterium. Hence, we assumed *β* = 2 ± 1 mg cellular water/mg CDW for *V. splendidus* sp. 1A01. The difference between *x* and *y* sets the upper bound on the weight of cellular water in a wet pellet, *xβ*_*max*_, which leads to 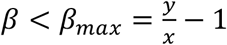. If the supernatant after washing was carefully removed, *β*_*max*_ is typically below 5. Within 0 < *β* < *β*_*max*_*∼*5, *α* is only weakly dependent on *β* (Fig. S1), and hence robust to error in the estimation of *β* within this range. This weak dependence results from the much lower density of the NaCl washing solution than the cellular mass density (*ρ*= 0.0306 for 0.52 M NaCl washing solution compared to *∼*0.5 mg CDW/mg cellular water).

Eq. 4 can also be rewritten as follows:

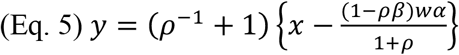

Eq. 5 predicts that a linear relation is obtained when *y* is plotted against *x*, and that *ρ* and *α* can be estimated from its slope and *x*-intercept, respectively. To test this prediction, we dispensed the culture grown at 1.25x SW on glucose into three aliquots of the same volume and obtained three washed wet pellets. Then we added back 0, 100, and 200 mL of the washing solution to each wet cell pellet and measured *x* and *y* for each pellet. As predicted, *x* and *y* showed a linear relation (Fig. S2). *ρ*= 2.98 × 10^−2^ was estimated from the slope, which is close to the measured value (*ρ*= 0.0306). *α* = 0.550 was estimated from the *x*-intercept with *β* = 2, which is also close to those estimated from Eq. 4 for the three pellets (*α* = 0.547 ± 0.001). This result also demonstrates that the direct estimate from Eq. 4 is precise enough.

After growing *V. splendidus* sp. 1A01 in a number of different media (in marine broth, and in minimal media on glucose, galactose, and glycerol) at different osmolarities (1x and 1.25x SW), we obtained (from Fig. S3) the following conversion factor *α* between CDW and OD_600_ (in units of mg CDW/OD_600_/mL), which varies linearly as a function of growth rate µ:

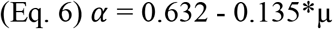

Thus, on glucose (µ ≈ 0.8 h^-1^), *α* is approximately 0.5 mg CDW/OD_600_/mL.

### Constructing a model of biomass composition

*V. splendidus* sp. 1A01 was grown in marine broth and in minimal medium on glucose, glucosamine, glycerol, and galactose. RNA and protein were quantified in each culture (in units of mg/OD_600_/mL) and plotted against growth rate, yielding the linear relations shown in Figs. S4 and S5. For the model, RNA and protein measurements corresponding to growth on glucose were used to build the biomass reaction (after converting from mg/OD_600_/mL to mg/mg CDW using Eq. 6). These measurements quantify the *total* RNA and protein in a biomass sample, not the relative proportions of the monomers that make up RNA and protein. As an approximation frequently made in the FBA literature^40^, the relative proportions of the 20 amino acids that make up protein were inferred from their relative proportions in the protein-coding regions of *V. splendidus* sp. 1A01’s genome. Likewise, the relative proportions of the 4 nucleotides that make up RNA were inferred from their relative proportions in the rRNA-coding regions of *V. splendidus* sp. 1A01’s genome (rRNA makes up the bulk of RNA in bacterial cells^46^). The osmolytes (glutamate and glutamine) were quantified at the standard osmolarity of 1x SW on glucose (10 mM), galactose (10 mM), glycerol (20 mM), and GlcNAc (10 mM). Like with RNA and protein, osmolyte measurements corresponding to growth on glucose (see Table S2) were used to build the model’s biomass reaction. The unquantified components of biomass (namely, DNA, LPS, lipids, murein, inorganic ions, and soluble pools) reflect the *E. coli* model iAF1260^47^. Because they represent 20% of *E. coli* biomass and only 17% of *V. splendidus* sp. 1A01, the total fraction of biomass these unquantified components represent was scaled down for *V. splendidus* sp. 1A01, while keeping their relative proportions the same as in *E. coli*. As with RNA and protein, the relative proportions of the 4 nucleotides that make up DNA were inferred from their relative proportions in *V. splendidus* sp. 1A01’s draft genome. Since populating parts of 1A01’s biomass reaction with coefficients from *E. coli* is an approximation, a sensitivity analysis of the model’s performance to slight deviations from *E. coli* (see Fig. S6) was carried out. In this analysis, for every unquantified component of biomass (DNA, LPS, lipids, murein, inorganic ions, or soluble pools), the fraction of biomass it represents was arbitrarily raised or lowered by up to 25%, which led to only negligible changes in flux through the biomass reaction (less than 1.4%). The complete biomass reaction is shown in Table S3, along with all calculations that lead from the above measurements and approximations to the final biomass reaction.

Although the model’s biomass reaction reflects the biomass composition of *V. splendidus* sp. 1A01 grown on glucose, in reality, the biomass composition varies based on growth rate (GR), as shown in Figs. S3 and S4. We therefore wrote a MATLAB script (Script S1) to incorporate 1A01’s GR-dependent biomass composition in FBA.

### Reconstruction and gap-filling of draft metabolic model

Estimated to be 99.4% complete by CheckM^48,49^, the annotated genome of *V. splendidus* sp. 1A01 was uploaded to BioCyc, where it was reconstructed into a draft metabolic model (in the form of an SBML file) using Pathway Tools^50^. The model’s reactions and metabolites therefore conform to BioCyc nomenclature (except where reactions from the BiGG^51^ database were inserted to fill gaps in metabolic pathways). The SBML file was then imported into MATLAB for model curation^40^, which included the following gap-filling. If *V. splendidus* sp. 1A01 grew on a given carbon source *in vivo* (growth being defined as achieving an OD_600_ increase of at least 0.9 after 20 hours), the minimum number of reactions were added to the model that enabled flux through its biomass reaction on the same carbon source. Exchange reactions were only added to the model for such experimentally verified carbon sources.

### Flux Balance Analysis

Flux Balance Analysis (FBA) is a widely adopted computational method to model cellular metabolism^18^. When applied to a GSMM, FBA predicts the steady-state distribution of fluxes that optimizes a certain objective function. In this work, to simulate bacterial growth, biomass production was chosen as the objective function. To calculate the NGAM, ATP production was chosen as the objective function (see *Estimation of GAM and NGAM*). Briefly, FBA is implemented as a linear programming problem and formulated in matrix notation as follows:

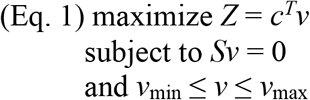

where *Z* denotes the objective function, *c* the relative weight of every reaction in the objective function, *v* the flux distribution, *S* the stoichiometric matrix, v_min_ the lower bounds on metabolic fluxes, and v_max_ the upper bounds. We performed FBA in MATLAB using the Gurobi optimizer^52^. A simple script for performing FBA with the *V. splendidus* sp. 1A01 model is provided (Script S2).

### Estimation of GAM and NGAM

To determine the GAM and NGAM of *V. splendidus* sp. 1A01, we measured batch-culture growth rate and net carbon influx on glucose and galactose (Fig. 4c). For a detailed description of growth medium, measurement and calculation of growth rate, as well as measurement and calculation of substrate intake and organic acid excretion, please see ref. 53. During constant exponential growth on glucose in batch culture, *V. splendidus* sp. 1A01 excreted acetate at a constant rate due to overflow metabolism^54^. The rate of acetate accumulation was subtracted from the rate of substrate depletion to calculate the net carbon influx (or carbon utilization rate). By plotting carbon utilization rate against growth rate and taking the y-intercept of the line (see Fig. 4c), we obtained the baseline rate of carbon utilization by *V. splendidus* sp. 1A01 in the absence of biomass production. We then imposed this carbon utilization rate in FBA (by manipulating upper and lower bounds on substrate uptake and acetate secretion fluxes) and maximized ATP production (while setting flux through the biomass reaction to zero) to arrive at the value of the NGAM, which, in the model, is its own ATP hydrolysis reaction (ATP + H_2_O → ADP + Pi). The GAM (which can be found in Table S3, as part of the model’s complete biomass reaction) was also calculated using FBA, by fitting the model’s performance to the line in Fig. 4c. More precisely, we imposed the observed carbon utilization rates in FBA (again, by manipulating upper and lower bounds on substrate uptake and acetate secretion fluxes) and increased the GAM until FBA-predicted growth rates matched experimental values.

### Model validation

A metabolic model must be validated against experimentally measured growth on substrates other than those used to parameterize the model (in this work, 5 mM glucose and 5 mM galactose). Therefore, batch cultures of *V. splendidus* sp. 1A01 were grown on 10 mM pyruvate and 5 mM N-acetyl-glucosamine (the monomer of chitin, on which particles 1A01 was first isolated^5^), while, again, tracking growth rate, substrate depletion, and acetate accumulation (Table S4). Experimentally measured rates of substrate uptake and acetate secretion were then used, in FBA, to constrain the corresponding fluxes in the model to their observed values, before optimizing flux through the biomass reaction, thereby obtaining theoretically optimal growth rates, which, for model validation, were compared to experimentally measured growth rates.

## Results

Due to scientific knowledge gaps with regards to protein function and gene-to-protein mapping, genome annotations are, in general, incomplete^55^. Since draft models are automatically reconstructed from genome annotations^40^, the shortcomings of genome annotations propagate directly to draft models, which display missing reactions in many metabolic pathways^39^. The process of restoring these missing reactions, and obtaining a functional metabolic model, is called gap-filling^39^, and demands phenotypic data^56^. In order to test its metabolic capabilities, *V. splendidus* sp. 1A01 was first cultured on 78 carbon sources for 10 days to obtain a coarse growth phenotype (see Fig. S7). Out of these 78, *V. splendidus* sp. 1A01 successfully grew on 35 carbon sources (Table 1a). After gap-filling the relevant metabolic pathways (see Methods), the model yielded growth on all 35 carbon sources (Table 1a). Out of the 43 carbon sources *V. splendidus* sp. 1A01 failed to grow on (Table 1b), 10 of them enabled growth according to the model (Table 1c), provided that, in the model, they freely diffused through the cell’s inner membrane (*i*.*e*. they did not require designated transporters to enter the cytoplasm). If, on the contrary, they were barred from freely diffusing through the cell’s inner membrane (*i*.*e*. if they required designated transporters to enter the cytoplasm), then only 4 substrates out of 43 enabled growth in the model (Table 1c). The discrepancy may be due to the lack of expression of these transporters in the conditions studied.

**Table 1.**
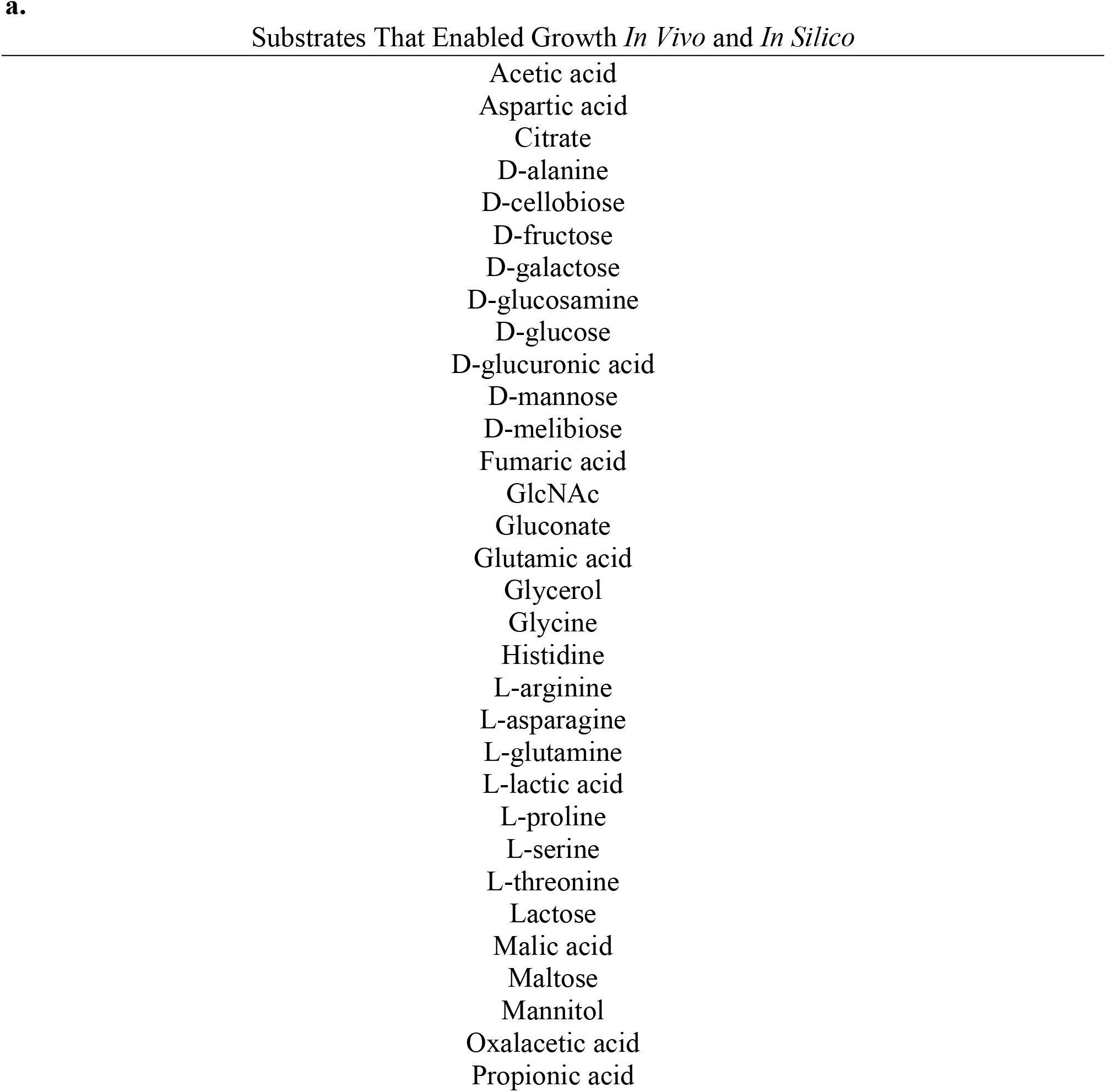
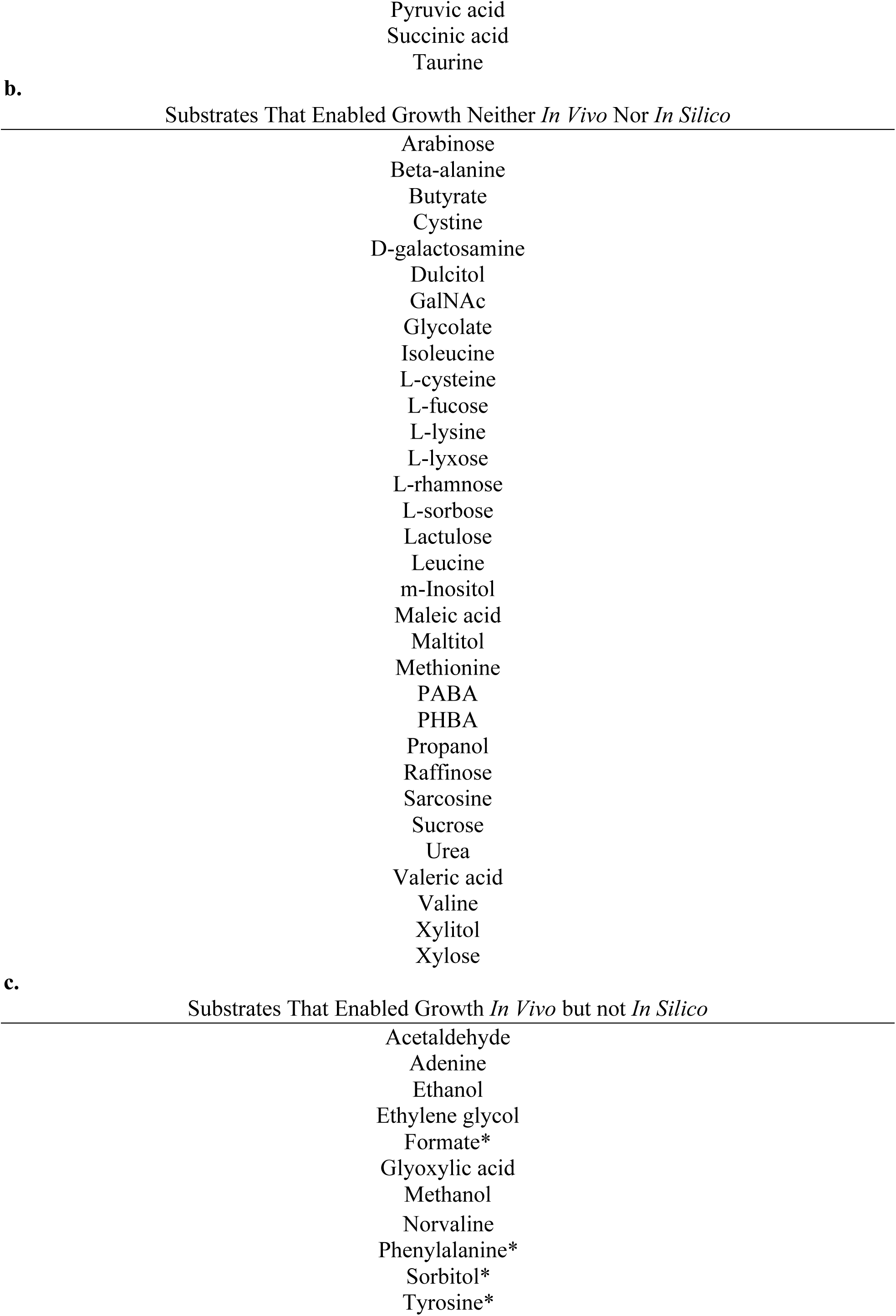
Experimental Phenotyping. **a**. The 35 carbon sources *V. splendidus* sp. 1A01 can grow on, both *in vivo* and *in silico*. **b**. The 43 carbon sources *V. splendidus* sp. 1A01 failed to grow on, both *in vivo* and *in silico*. **c**. The 10 carbon sources *V. splendidus* sp. 1A01 grew on *in vivo* but not *in silico*. The 4 carbon sources that enabled growth *in silico* when diffusing freely into the periplasm but no further are indicated by asterisks. Of these 4 carbon sources, sorbitol is the only one that did not yield growth when diffusing directly into the cytoplasm (without active transport) because its metabolism requires a phosphorylation reaction catalyzed by a transporter.

Following gap-filling, the model contained 1867 reactions and 1565 metabolites (Table 2). To further characterize the model, every reaction was assigned, if possible, to a metabolic pathway in BioCyc, and every metabolic pathway to a broad functional category. Fig. S8 shows the relative distribution of these broad functional categories across all assigned reactions.

**Table 2.**
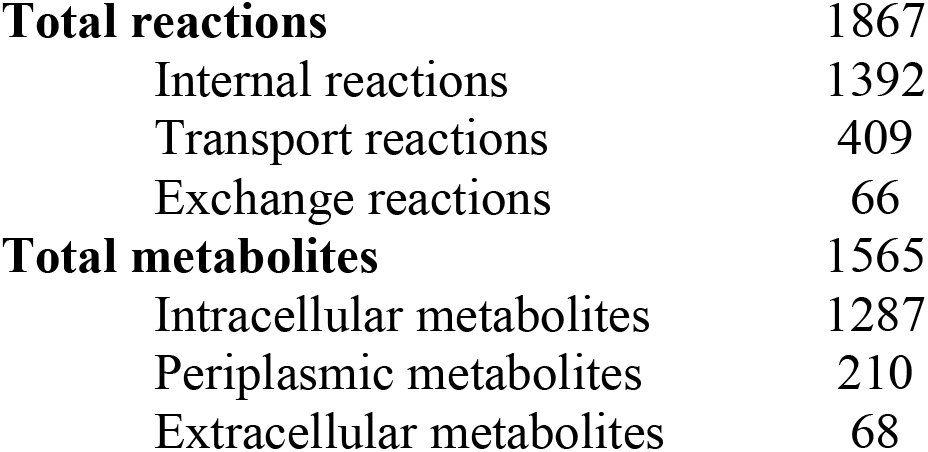
Overview of *V. splendidus* sp. 1A01 Model.

|  |  |
| --- | --- |
| **Total reactions** | 1867 |
| Internal reactions | 1392 |
| Transport reactions | 409 |
| Exchange reactions | 66 |
| **Total metabolites** | 1565 |
| Intracellular metabolites | 1287 |
| Periplasmic metabolites | 210 |
| Extracellular metabolites | 68 |

While enzyme-catalyzed reactions make up the bulk of a metabolic model, it contains also a biomass reaction that converts cellular building blocks (such as amino acids and nucleotides) into biomass, in experimentally measured proportions^40^. Quantifying these proportions is a vital part of model curation. Instead of measuring the exact concentration of every chemical compound to be found in biomass, it is both sufficient (for modeling purposes) and convenient (experiment-wise) to measure the overall macromolecular composition, and fill the leftover knowledge gaps with data from the better-studied *E. coli*, which, like *V. splendidus* sp. 1A01, is a Gram-negative gamma proteobacterium^57^. With regards to the biomass coefficients supplied by *E. coli*, a sensitivity analysis (described in Methods) was carried out and showed the model’s performance to be robust against slight deviations from the exact values in *E. coli* (see Fig. S6).

Protein, RNA, and osmolytes were expected to dominate the macromolecular composition of *V. splendidus* sp. 1A01, and were quantified accordingly. The contribution of RNA and proteins were measured in culture samples of *V. splendidus* sp. 1A01 grown on a variety of carbon sources, covering a spectrum of growth rates (Figs. S4 and S5). However, raw measurements of RNA and protein are in units of mg/OD_600_/mL, which must be converted to mg/mg CDW to become biologically meaningful. The conversion factor between OD_600_*mL and mg CDW was obtained by measuring the CDW of culture samples (of known OD_600_) across a range of growth rates (Fig. S3). As shown in Fig. 2, the RNA and protein composition of *V. splendidus* sp. 1A01, together with the strain’s CDW, all vary linearly as a function of growth rate. Together, they define the GR-dependent biomass composition of *V. splendidus* sp. 1A01.

**Fig. 2.**
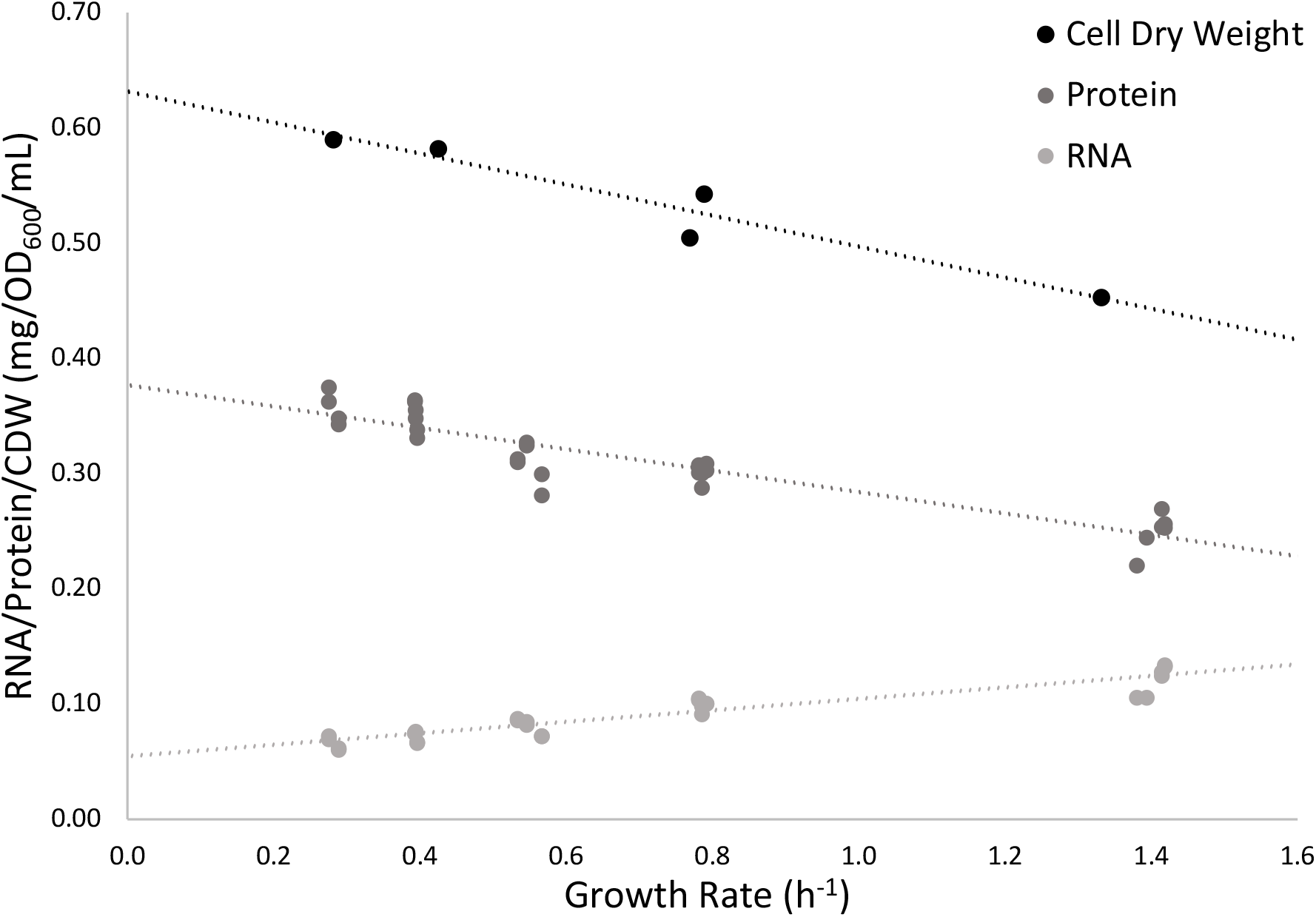
The Growth-Rate Dependence of RNA, Protein, and Cell Dry Weight. *V. splendidus* sp. 1A01 was grown in marine broth and in minimal medium on glucose, GlcN, glycerol, and galactose.

The biomass composition of *V. splendidus* sp. 1A01 growing on glucose (at a growth rate of 0.79 h^-1^) is shown in Fig. 3a. (In Fig. S9 is shown the biomass composition of *E. coli*, which is qualitatively similar.) For comparison, Fig. 3b shows the biomass composition of *V. splendidus* sp. 1A01 growing on galactose, at the lower growth rate of 0.29 h^-1^. As expected from Fig. 2, due to the higher growth rate associated with glucose compared to galactose, RNA accounts for a visibly greater percentage of CDW in Fig. 3a than in Fig. 3b. We selected the biomass composition of *V. splendidus* sp. 1A01 growing on glucose to build the model’s biomass reaction (which can be found in Table S3).

**Figure 3.**
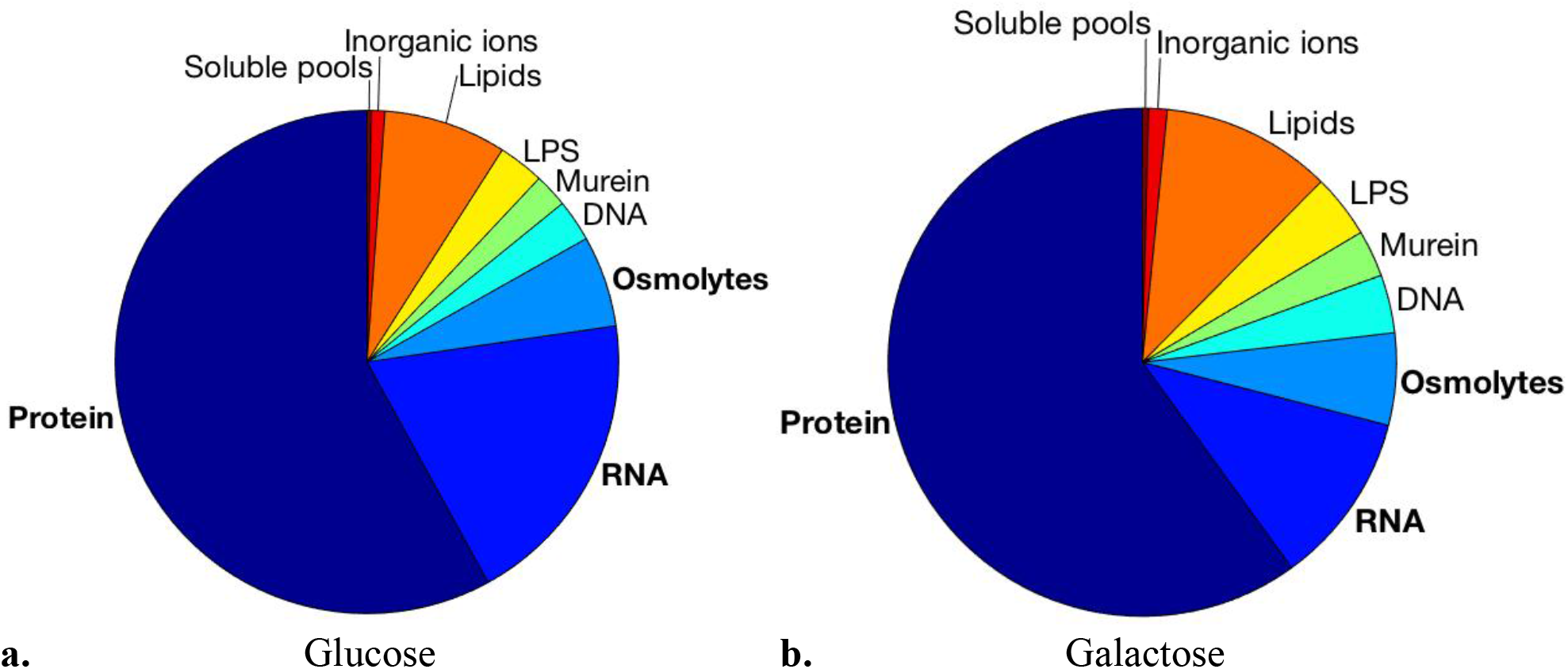
Macromolecular Biomass Composition of *V. splendidus* sp. 1A01 Growing on Glucose (a.) and Galactose (b.) In bold are shown the three components of biomass (protein, RNA, and osmolytes) for which experimental data was collected. Together, they make up 83% of the CDW of *V. splendidus* sp. 1A01 (on glucose). The remaining 17% of its biomass (DNA, LPS, lipids, murein, inorganic ions, and soluble pools) reflect the *E. coli* model iAF1260^47^.

The third and final step of model curation is to estimate the NGAM, which corresponds to the amount of energy the cell consumes just to survive^40^ (by, for example, maintaining the integrity of its membrane), and the GAM, which corresponds to the amount of energy the cell consumes to produce biomass^40^ (by, for example, polymerizing amino acids into protein). To estimate these values, *V. splendidus* sp. 1A01 was grown in batch culture on glucose and on galactose as the sole carbon substrate. The optical density, concentration of the carbon substrate, and concentration of the excreted acetate were measured at various time intervals throughout exponential batch-culture growth. From the temporal dependence of the optical density measurements, growth rates were deduced (0.79 h^-1^ for glucose and 0.29 h^-1^ for galactose). Plotting the concentration of the carbon substrate or the excreted acetate against optical densities, we obtained the consumption yields of growth on glucose and on galactose, together with the excretion yield of acetate, as the slopes of the respective plots (Fig. 4ab). The difference of the consumption and excretion yields (in units of carbon monomer per OD_600_) was then multiplied by the growth rate to obtain the carbon utilization rate (*i*.*e*. the rate at which carbon monomers are either incorporated into biomass or used for energy biogenesis) in each growth medium. The resulting carbon utilization rates were plotted in Fig. 4c against the corresponding growth rates. The y-intercept of this plot, 4.22 mM-C/OD_600_/h, represents the rate of carbon utilization by *V. splendidus* sp. 1A01 at zero growth rate, when carbon is utilized, not to produce biomass, but strictly to generate energy for cell “maintenance.” Using FBA, we found the maximal rate of ATP production given this carbon utilization rate and zero flux through the biomass reaction. The NGAM corresponds to this maximal rate of ATP production in the absence of growth, and shows up in the model as the minimum allowable flux (12.8 mmol/gCDW/h) through an ATP hydrolysis reaction, which must be satisfied under all conditions^40^. However, the GAM (measured at 15.8 mmol/gCDW) features in the model’s biomass reaction^40^, and reflects the slope of the line in Fig. 4c (23.3 mM-C/OD_600_). For comparison, the *E. coli* model iAF1260^47^ has an NGAM of 8.39 mmol/gCDW/h (vs. 12.8 mmol/gCDW/h in 1A01) and a GAM of 59.81 mmol/gCDW (vs. 15.8 mmol/gCDW in 1A01).

As mentioned above, we selected the biomass composition of *V. splendidus* sp. 1A01 grown on glucose to build the model’s biomass reaction (Table S3). However, as shown in Figs. 3 and 4, 1A01’s biomass composition varies based on growth rate. Therefore, after quantifying the GAM and NGAM, we investigated whether taking the GR-dependence of 1A01’s biomass composition into account (instead of applying the same glucose-derived biomass reaction to all conditions) would significantly improve the model’s predicted growth rates on carbon sources other than glucose. To wrap the GR-dependence of 1A01’s biomass composition into FBA (see Script S1 for full code), a growth rate on a given carbon source is first guessed, and the corresponding biomass composition incorporated into the model via the biomass reaction. FBA then uses the model to predict an optimal growth rate. If this optimal growth rate matches the initially guessed growth rate, the script stops. Otherwise, the cycle repeats itself, with the FBA-predicted growth rate generating a new biomass composition (and, by extension, a new biomass reaction in the model), until the script converges (*i*.*e*. until the input and output growth rates match). We found that the script converges to approximately the same final growth rate, regardless of the initially guessed growth rate. For example, on galactose, it converges to a growth rate of about 0.27 h^-1^, regardless of whether one initially guesses 0.01 h^-1^ or 1 h^-1^. Comparing the results of GR-dependent FBA to “standard” FBA with a glucose-derived biomass reaction, both applied to growth on galactose, we found that the two gave very similar growth rates (Fig. S10a). The two methods gave larger differences in fluxes, with about half of the fluxes showing differences of at least 20% between glucose and galactose (Fig. S10b), reflecting substantially different dry mass compositions. Overall, even though the GR-dependent biomass composition can be accommodated, running FBA on the glucose-derived model provides reasonable approximations of both growth rates and flux distributions.

**Figure 4.**
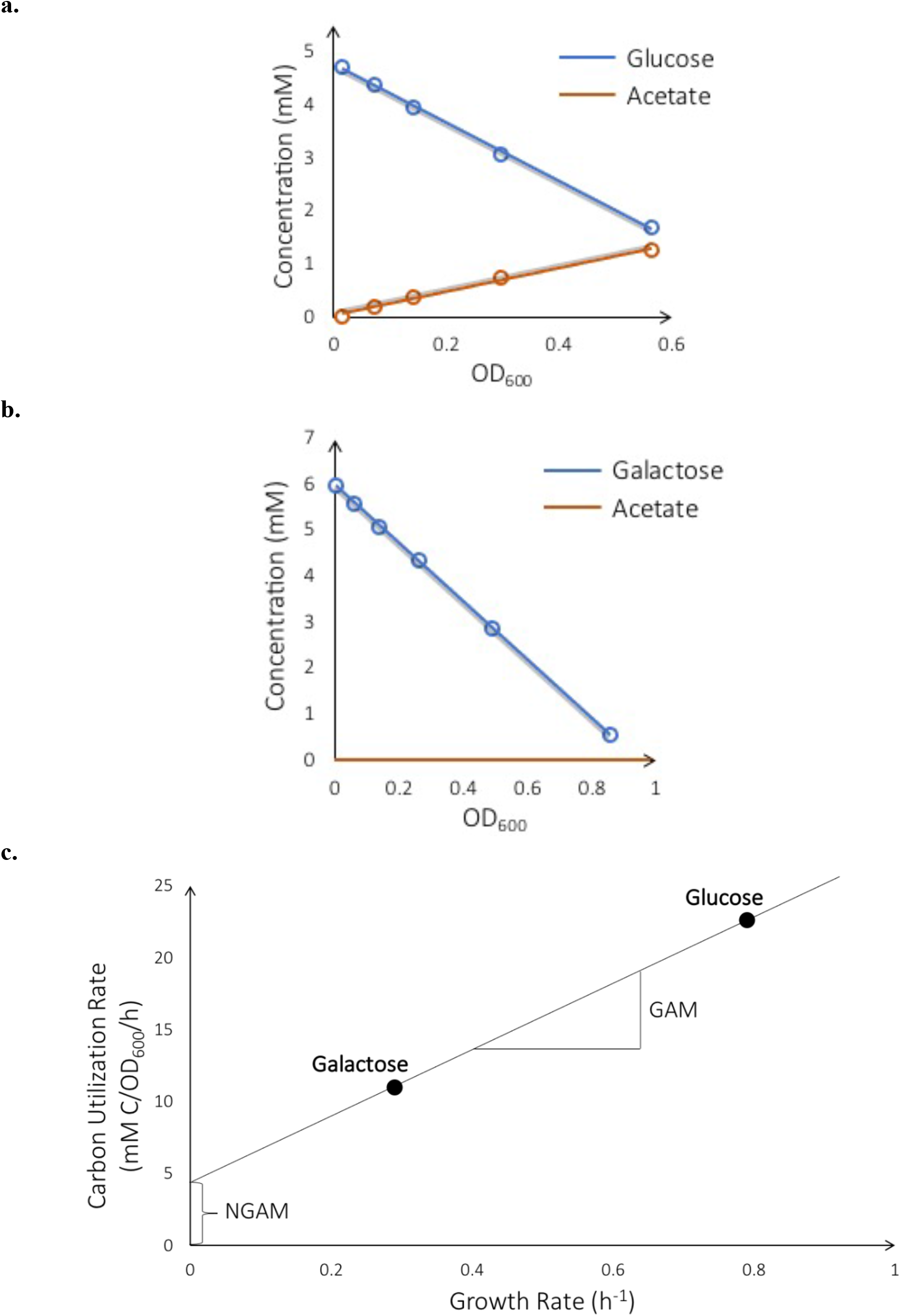
GAM and NGAM. **a.b**. Shown in red is acetate accumulation, in blue, glucose or galactose depletion. No acetate is secreted when *V. splendidus* sp. 1A01 is grown on galactose. The consumption yields of growth on glucose and galactose, together with the excretion yields of acetate, correspond to the slopes of blue and red plots respectively. **c**. The carbon utilization rate then corresponds to the difference between consumption and excretion yields, multiplied by the growth rate on each carbon source. The GAM is calculated based on the slope of the line, the NGAM based on its y-intercept. Regarding the units, mM C corresponds to mM of carbon atoms in the substrates.

Next, the metabolic model was quantitatively tested against growth on carbon substrates other than those used to parameterize the model (*i*.*e*. glucose and galactose). This time, *V. splendidus* sp. 1A01 was grown in batch on pyruvate and on N-acetyl-glucosamine. Again, the optical density, concentration of substrate, and concentration of acetate were measured at time intervals throughout exponential batch-culture growth (Fig. 5ab). Rates of substrate consumption and acetate secretion were measured as before, only now they served as input into FBA rather than for inferring the parameters of the model (namely, the GAM and NGAM). More precisely, as depicted in Fig. 5c, the model’s carbon uptake flux (v_carb_) and acetate secretion flux (v_ace_) were set to the observed values, and flux through the biomass reaction (v_bio_) was optimized using FBA. For comparison, the same process was repeated for glucose and galactose. The good agreement between experiment and theory we observe for these last two substrates (as shown in Fig. 5d) is expected, given that the model was trained on them. However, we see the same agreement for pyruvate and, to a less extent, N-acetyl-glucosamine. (This imperfect agreement, in the case of N-acetyl-glucosamine, might be due to the coupling of carbon and nitrogen sources in this compound^58^.) We conclude that the model’s predictive power extends beyond merely its training data.

**Figure 5.**
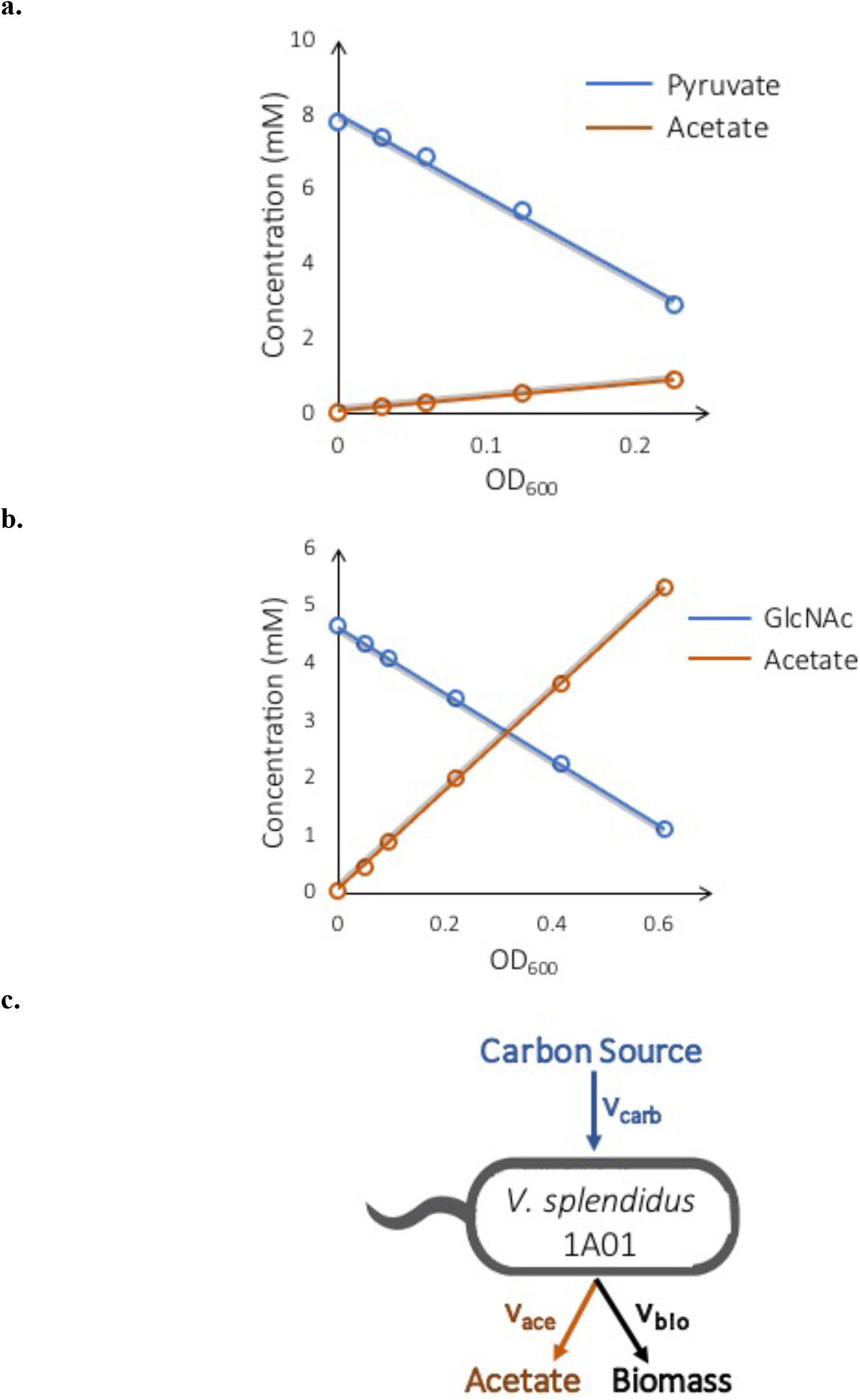
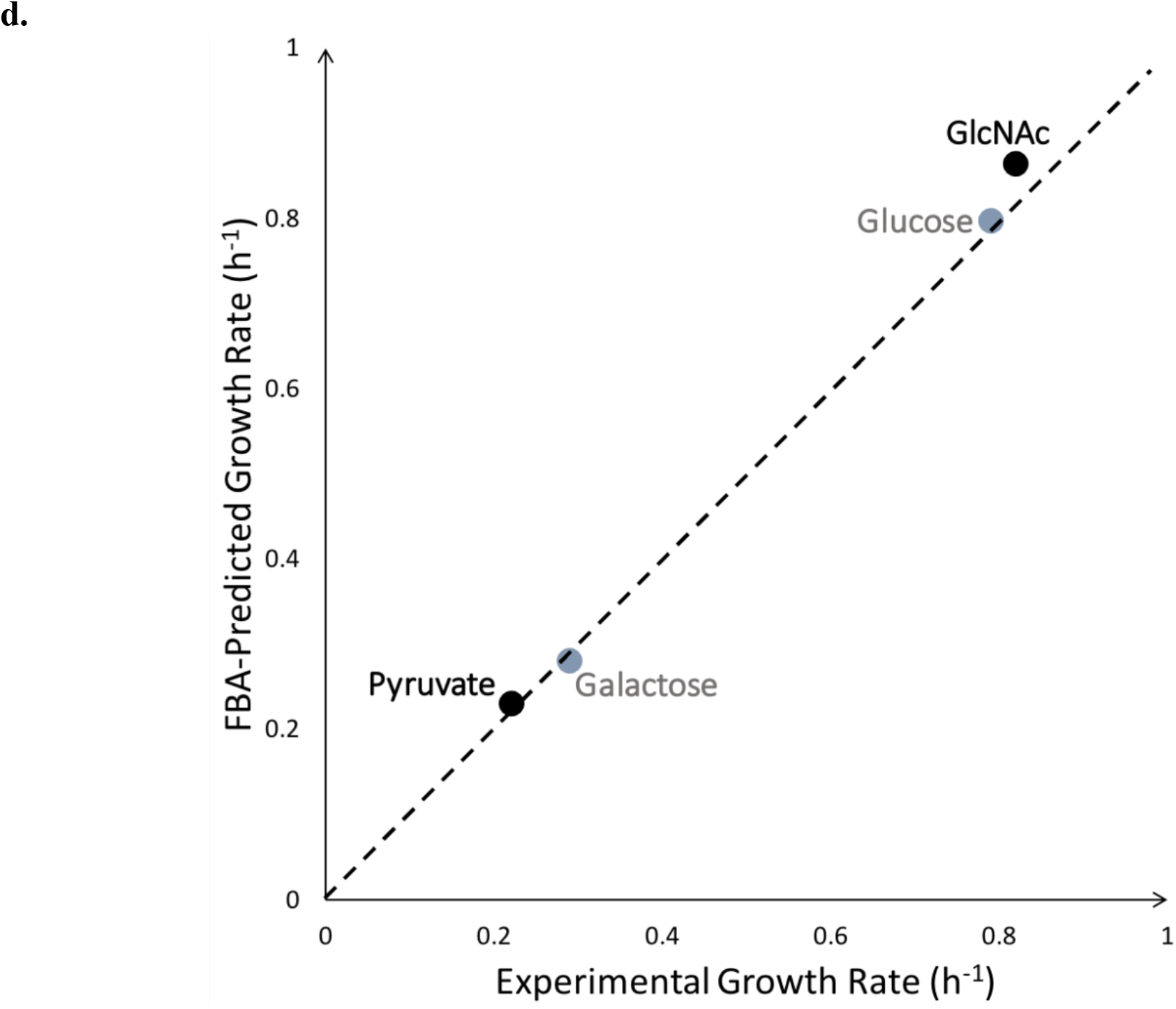
Quantitative Testing of Model Predictions. **a.b**. Shown in red is acetate accumulation, in blue, pyruvate or N-acetyl-glucosamine (GlcNAc) depletion. **c**. v_bio_ denotes flux through the biomass reaction, while v_carb_ and v_ace_ denote carbon uptake flux and acetate secretion flux, respectively. **d**. Along the x-axis are plotted experimentally measured growth rates of *V. splendidus* sp. 1A01, and, along the y-axis, theoretical growth rates predicted by the model. Shown in grey are substrates the model was trained on (glucose and galactose). Shown in black are substrates the model is being tested against (pyruvate and N-acetyl-glucosamine). The dashed line denotes perfect agreement between model and experiment. Note the model cannot be fit arbitrarily well to the training data, which is why the two points for glucose and galactose do not lie perfectly along the dashed line.

## Discussion

In this work, a GSMM was reconstructed for *V. splendidus* sp. 1A01, a chitin-degrading opportunistic pathogen in the ocean^5,9^. The model reconstruction required measuring a number of physiological parameters, which gave rise to the following technical challenges, partly due to the high salt concentration found in seawater (and in synthetic culture media, like MBL, that simulate seawater).

First, the GAM and NGAM are conventionally found by growing microbes in a chemostat^40^. However, adapting wild organisms to chemostat growth is often difficult, and we opted to grow batch cultures of *V. splendidus* sp. 1A01 on different carbon sources, at the maximal growth rate permitted by each carbon source (*i*.*e*. at saturating concentrations of the substrate). Because different carbon sources allow for different maximal growth rates, we were able to obtain a spread of growth rates that is obtained in a chemostat by tuning the dilution rate on a single carbon source. However, because strong overflow metabolism can occur near a strain’s optimal growth rate^54^, acetate excretion also had to be measured. We regard this method of measuring the GAM and NGAM as equivalent in accuracy to running a chemostat^40^.

Second, marine microbes differ in their biomass composition from other bacterial strains, due to the high salt concentration in seawater. All bacteria must produce osmolytes to balance external osmolarity, but seawater is of greater salinity than M9^59^, the traditional culture medium of *E. coli*. As a result, marine bacteria must produce more osmolytes. We experimentally measured the percentage of 1A01’s cell dry weight that osmolytes represent, in order to arrive at a more accurate biomass reaction in the model.

Third, the high medium osmolarity made it challenging to measure the conversion factor of OD_600_ to CDW, which must be measured to convert experimental measurements (often in units of OD_600_) to the unit of flux in FBA, mmol/gCDW/h, with gCDW corresponding to grams of CDW. To measure this conversion factor in *E. coli*, the first step is to spin down a culture sample in a centrifuge and resuspend it in water, in order to wash away extracellular metabolites contained in the medium. However, if *V. splendidus* sp. 1A01 is resuspended in water, cells burst due to the larger osmotic shock, leading to significant loss of biomass. We therefore had to develop a novel experimental technique (described in Methods) to measure the CDW of a marine bacterium like *V. splendidus* sp. 1A01.

In conclusion, we expect many of the technical issues we faced in building a model for *V. splendidus* sp. 1A01 to reappear when building models for other marine microbes. Thus, we hope this work will not only shed light on the metabolic capabilities and behavior of *V. splendidus* sp. 1A01, but will guide the reconstruction of GSMMs for the myriad other bacteria that populate our oceans.

## Software Availability

The genome-scale metabolic model of *V. splendidus* sp. 1A01, as well as the code to run FBA (Script S2) and GR-dependent FBA (Script S1) on this model, are available in a GitHub repository at https://github.com/ArionIfflandStettner/VibrioSplendidus1A01.

## Acknowledgments

This work was supported by the Simons Foundation through the PriME (Principles of Microbial Ecosystems) collaboration (Grant no. 542381 to SB, 542387 to TH, and 542395 to OXC).

We are grateful to Julia Schwartzman for estimating the completeness of the *V. splendidus* sp. 1A01 draft genome using CheckM^48,49^.

