## Supplementary Material for "A Genome-Scale Metabolic Model of Marine Heterotroph *Vibrio splendidus* sp. 1A01"

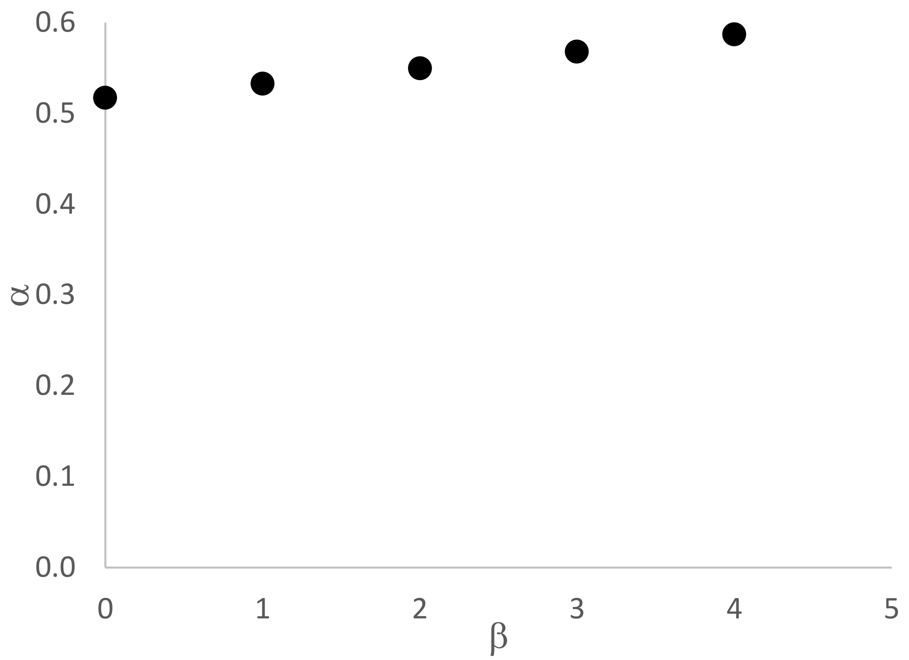


**Fig. S1. An example of CDW per $OD_{600}$·mL, $\alpha$, as a function of cellular water weight per CDW, $\boldsymbol{\beta}$.** The culture was grown in a minimal medium supplemented with 10 mM glucose at 1.25x SW. It can be seen that $\alpha$ is only weakly dependent on $\beta$. With $\beta = 2 \pm 1$ mg cellular water/mg CDW, $\alpha$ is estimated to be $0.536 \pm 0.015$ mg per $OD_{600}$·mL.

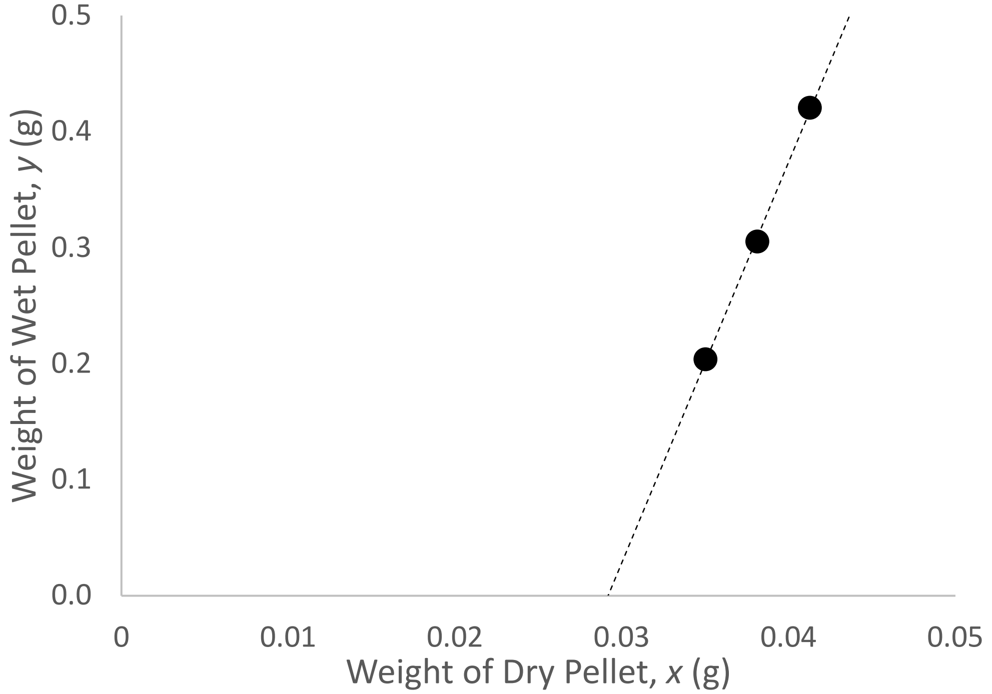


**Fig. S2. The linear relation between the weights of wet and dry cell pellets.** Cells were grown in minimal media at 1.25x SW supplemented with 10 mM glucose.

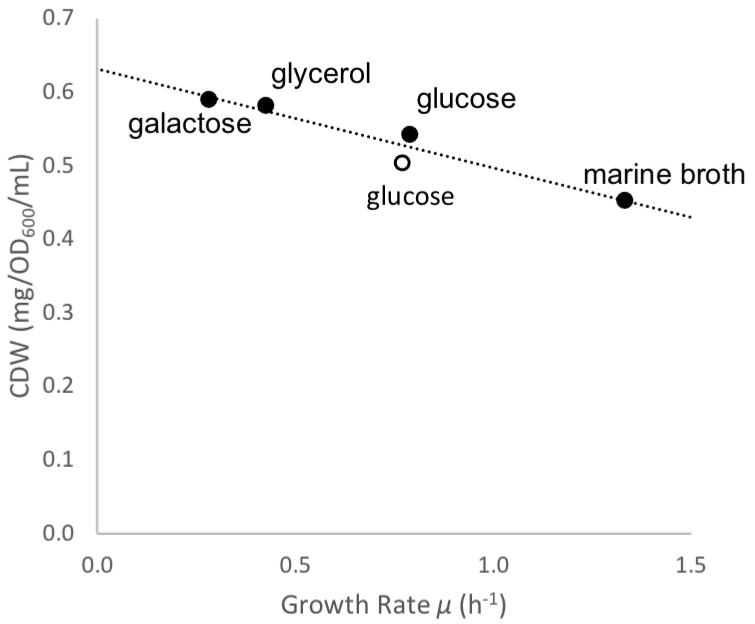


**Fig. S3. The growth-rate dependence of $\alpha$ with $\beta = 2$.** The osmolarities of the media are 1x SW (closed circles) and 1.25x SW (open circle). The concentrations of carbon sources in the minimal media are 10 mM for both glucose and galactose, and 20 mM for glycerol. For FBA analysis, $\alpha$ is estimated as a linear function (of the growth rate $\mu$) obtained from this plot ($\alpha = 0.632 - 0.135 \cdot \mu$).

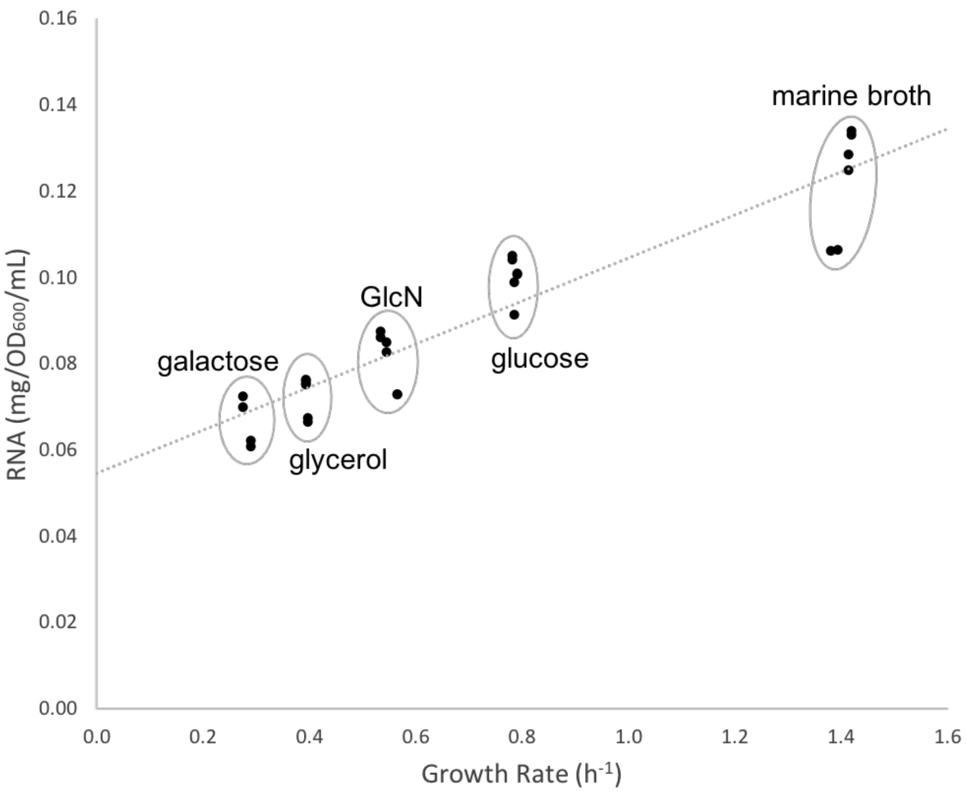


**Fig. S4. The growth-rate dependence of RNA production.** *V. splendidus* sp. 1A01 was grown in marine broth and in minimal media on glucose, GlcN, glycerol, and galactose.

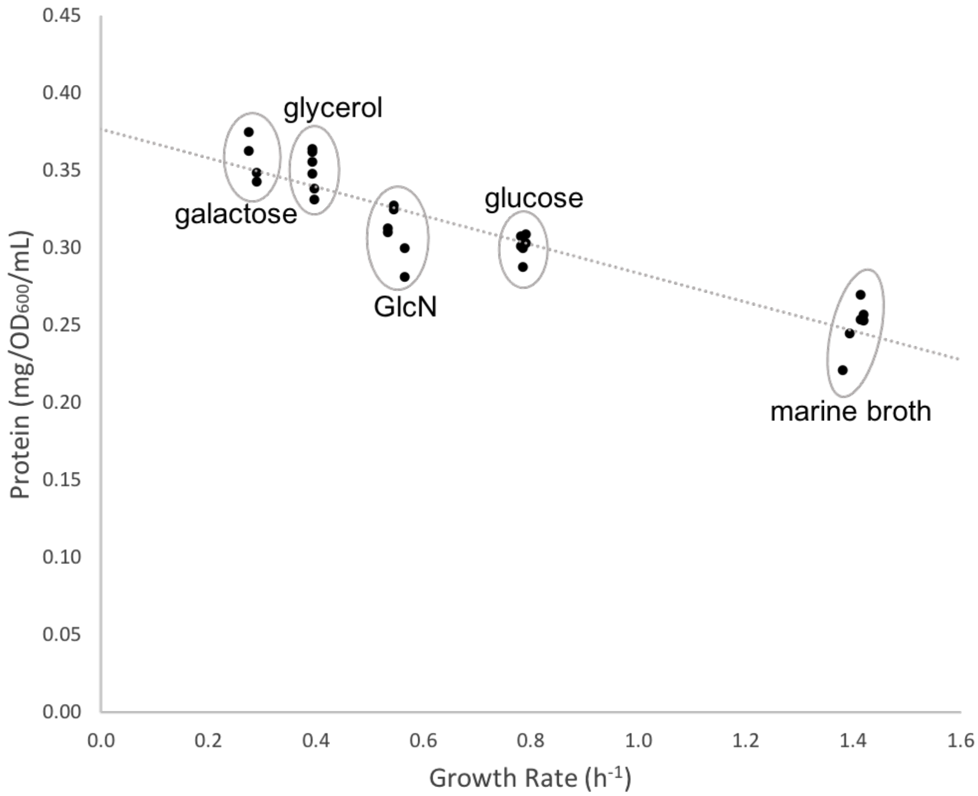


**Fig. S5. The growth-rate dependence of protein production.** *V. splendidus* sp. 1A01 was grown in marine broth and in minimal media on glucose, GlcN, glycerol, and galactose.

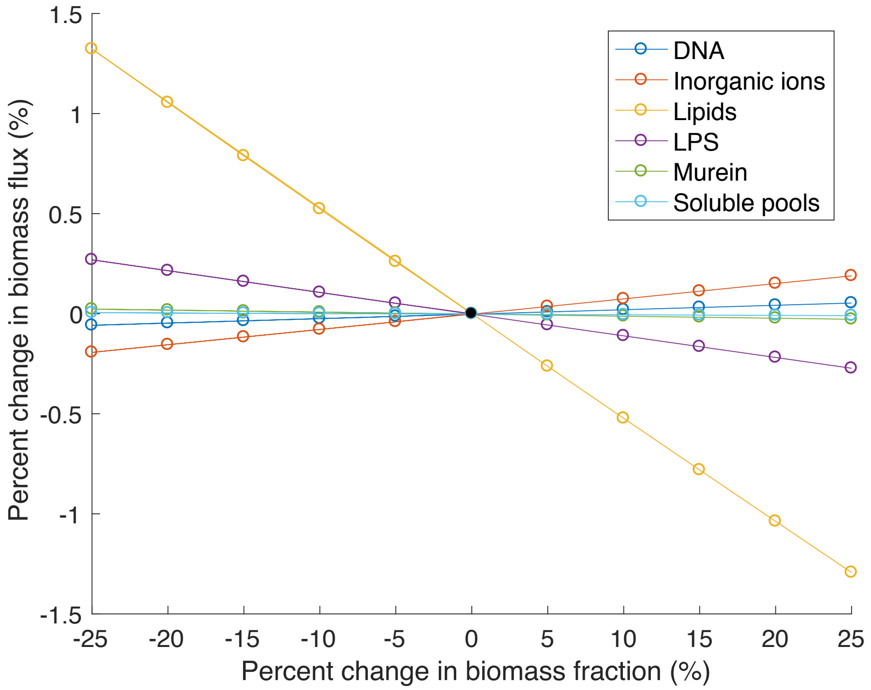


**Fig. S6. Sensitivity analysis to deviations from *E. coli* in the biomass reaction of the *V. splendidus* sp. 1A01 model.**

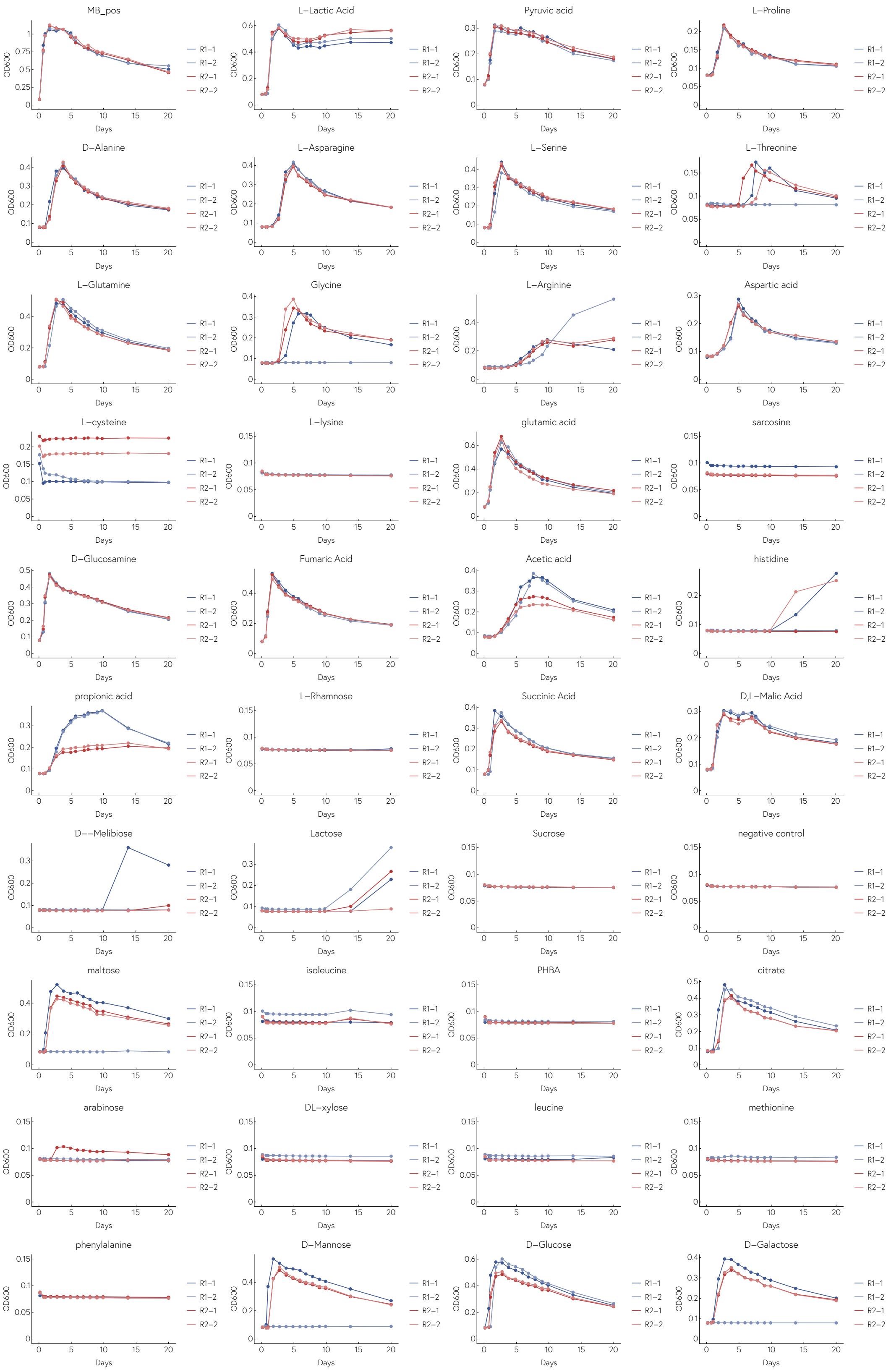

MB_pos
L−Lactic Acid
Pyruvic acid
L−Proline
D−Alanine
L−Asparagine
L−Serine
L−Threonine
L−Glutamine
Glycine
L−Arginine
Aspartic acid
L−cysteine
L−lysine
glutamic acid
sarcosine
D−Glucosamine
Fumaric Acid
Acetic acid
histidine
propionic acid
L−Rhamnose
Succinic Acid
D,L−Malic Acid
D−−Melibiose
Lactose
Sucrose
negative control
maltose
isoleucine
PHBA
citrate
arabinose
DL−xylose
leucine
methionine
phenylalanine
D−Mannose
D−Glucose
D−Galactose
OD600
Days
R1−1
R1−2
R2−1
R2−2

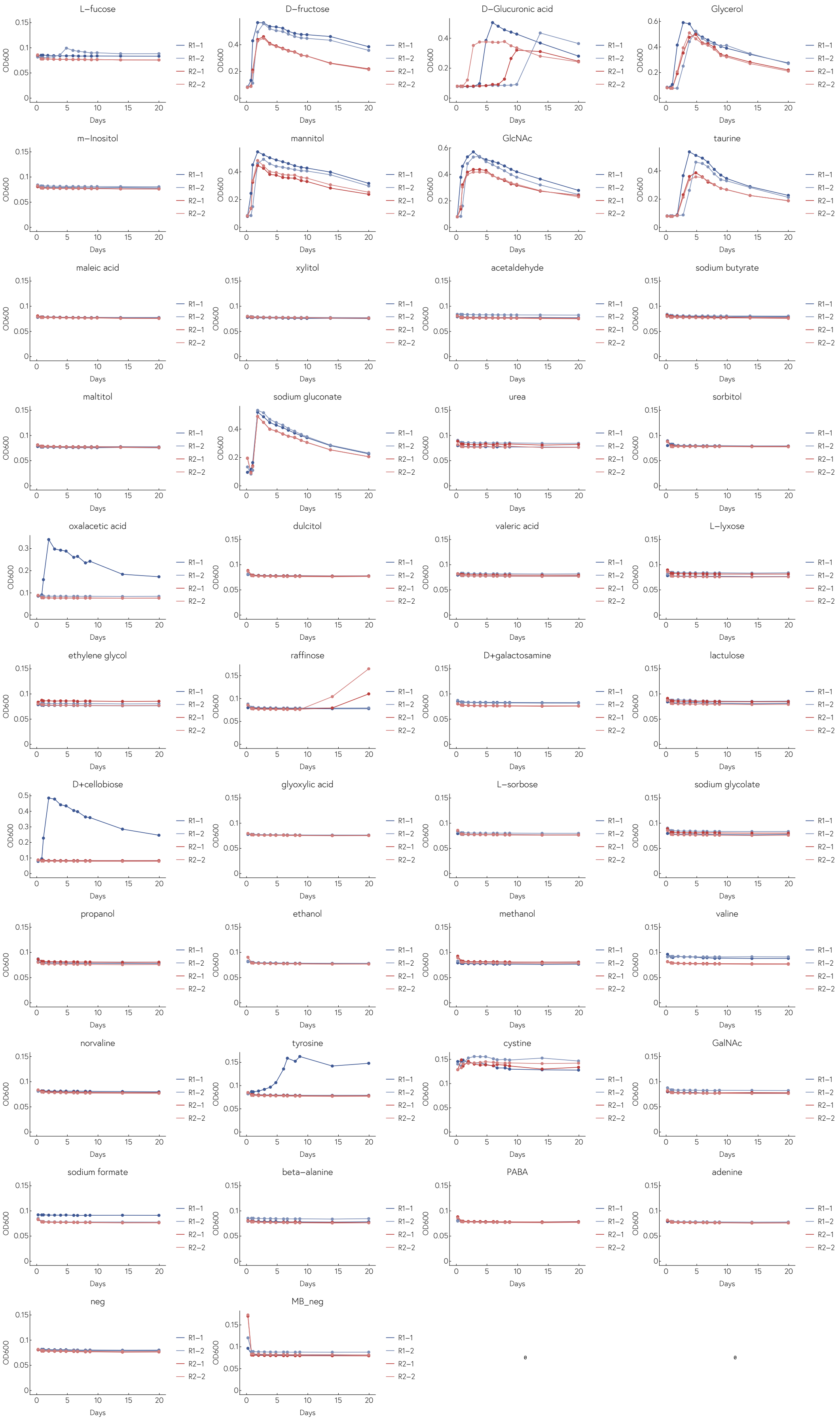
L−fucose
D−fructose
D−Glucuronic acid
Glycerol
m−Inositol
mannitol
GlcNAc
taurine
maleic acid
xylitol
acetaldehyde
sodium butyrate
maltitol
sodium gluconate
urea
sorbitol
oxalacetic acid
dulcitol
valeric acid
L−lyxose
ethylene glycol
raffinose
D+galactosamine
lactulose
D+cellobiose
glyoxylic acid
L−sorbose
sodium glycolate
propanol
ethanol
methanol
valine
norvaline
tyrosine
cystine
GalNAc
sodium formate
beta−alanine
PABA
adenine
neg
MB_neg
OD600
Days
R1−1
R1−2
R2−1
R2−2

**Fig. S7. Experimental phenotyping.** The full growth curves of *V. splendidus* sp. 1A01 on all 78 carbon sources. The replicates are technical (same original culture, separate wells) and biological (separate original cultures).

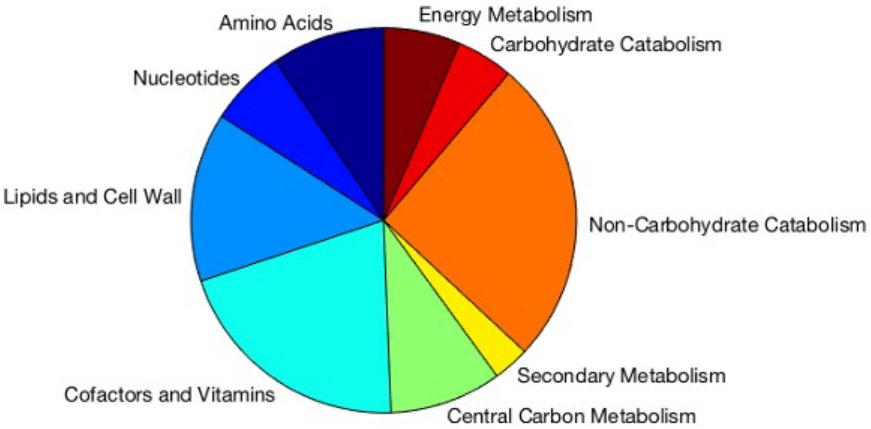


**Fig. S8. Distribution of pathway categories in *V. splendidus* sp. 1A01 model.** The different shades of blue and purple on the left side of the pie chart correspond to biosynthesis categories (*e.g.* biosynthesis of nucleotides).

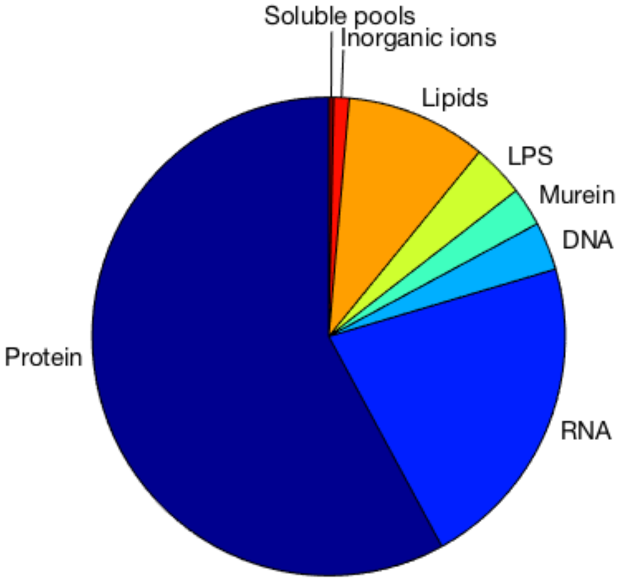


**Fig. S9. Macromolecular biomass composition of *E. coli* iAF1260.**

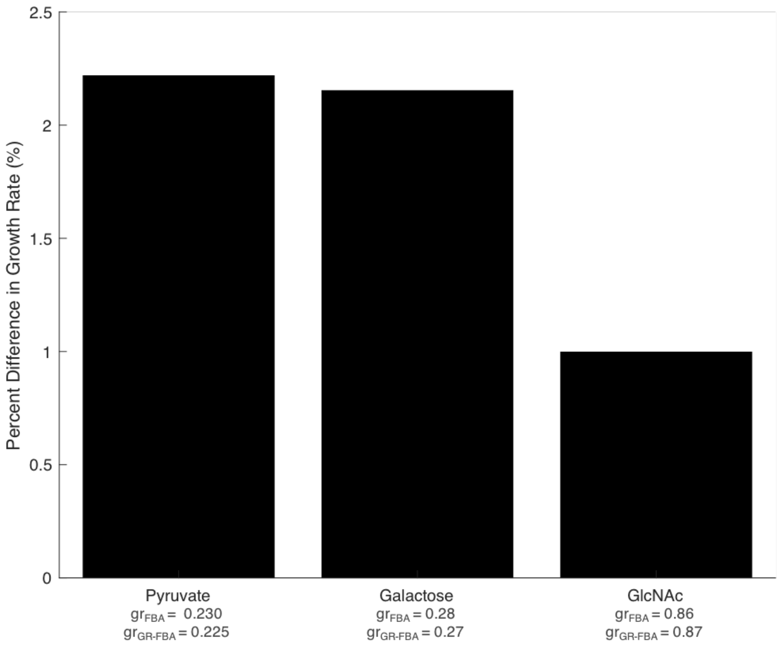


**a.**

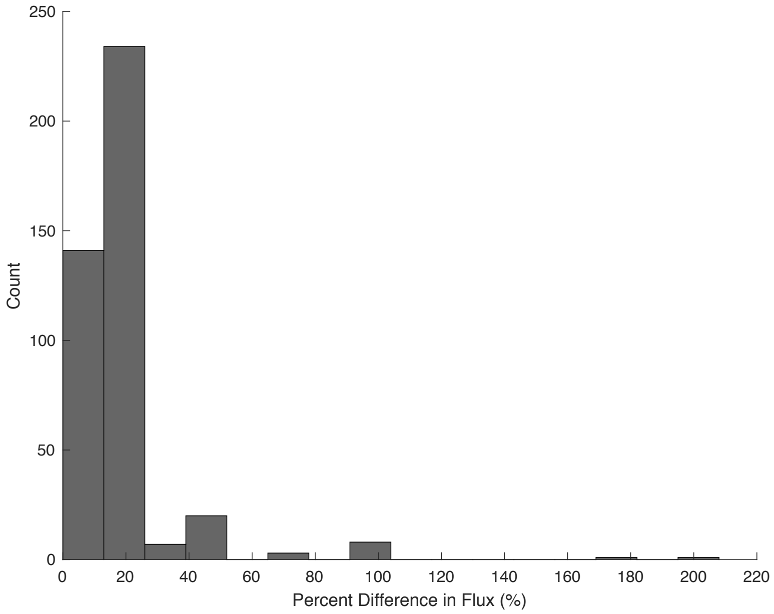


**b.**

**Fig. S10. Here we compare the output of FBA using the "standard" glucose-derived biomass reaction ($gr_{FBA}$) and the GR-dependent biomass reaction ($gr_{GR\text{-}FBA}$), when simulating growth of *V. splendidus* sp. 1A01 on various carbon sources. a.** Bar plot of the percent difference between the growth rates output by the two FBA methods, for growth on pyruvate, galactose, and N-acetylglucosamine (GlcNAc). Note that the deviation between the two methods is the smallest for GlcNAc, since the growth rates on GlcNAc and glucose are similar and so the biomass compositions are similar. **b.** Histogram of the percent difference between the fluxes output by FBA using the "standard" glucose-derived biomass reaction and the fluxes output by FBA using the GR-dependent biomass reaction, when simulating growth of *V. splendidus* sp. 1A01 on galactose.

**Table S1: Carbon Sources for Growth Phenotype Screening**

| Carbon Source | MW | Number of C Atoms | Concentration (mM) | Chemical Formula |
|---|---|---|---|---|
| L-Lactic Acid | 90.078 | 3 | 13.33 | C3H6O3 |
| Pyruvic Acid | 88.06 | 3 | 13.33 | C3H4O3 |
| L-Proline | 115.132 | 5 | 8.00 | C5H9NO2 |
| D-Alanine | 89.094 | 3 | 13.33 | C3H7NO2 |
| L-Asparagine | 132.119 | 4 | 10.00 | C4H8N2O3 |
| L-Serine | 105.093 | 3 | 13.33 | C3H7NO3 |
| L-Threonine | 119.12 | 4 | 10.00 | C4H9NO3 |
| L-Glutamine | 146.146 | 5 | 8.00 | C5H10N2O3 |
| Glycine | 75.067 | 2 | 20.00 | C2H5NO2 |
| L-Arginine | 174.204 | 6 | 6.67 | C6H14N4O2 |
| Aspartic Acid | 133 | 4 | 10.00 | C4H7NO4 |
| L-Cysteine | 121.15 | 3 | 13.33 | C3H7NO2S |
| L-Lysine | 146.19 | 6 | 6.67 | C6H14N2O2 |
| Glutamic Acid | 147.1 | 5 | 8.00 | C5H9NO4 |
| Sarcosine | 89.094 | 3 | 13.33 | C3H7NO2 |
| D-Glucosamine | 179.172 | 6 | 6.67 | C6H13NO5 |
| Fumaric Acid | 116.072 | 4 | 10.00 | C4H4O4 |
| Acetic Acid | 60.052 | 2 | 20.00 | C2H4O2 |
| Histidine | 155 | 6 | 6.67 | C6H9N3O2 |
| Propionic Acid | 74.079 | 3 | 13.33 | C3H6O2 |
| L-Rhamnose | 164.157 | 6 | 6.67 | C6H12O5 |
| Succinic Acid | 118.088 | 4 | 10.00 | C4H6O4 |
| D,L-Malic Acid | 134.087 | 4 | 10.00 | C4H6O5 |
| D-Melibiose | 342.297 | 12 | 3.33 | C12H22O11 |
| Lactose | 342.297 | 12 | 3.33 | C12H22O11 |
| Sucrose | 342.3 | 12 | 3.33 | C12H22O11 |
| Maltose | 342.297 | 12 | 3.33 | C12H22O11 |
| Isoleucine | 131.2 | 6 | 6.67 | C6H13NO2 |
| 4-Hydroxybenzoic Acid | 138.121 | 7 | 5.71 | C7H6O3 |
| Citric Acid | 192.123 | 6 | 6.67 | C6H8O7 |
| Arabinose | 150.13 | 5 | 8.00 | C5H10O5 |
| DL-Xylose | 150.13 | 5 | 8.00 | C5H10O5 |
| Leucine | 131.2 | 6 | 6.67 | C6H13NO2 |
| Methionine | 149 | 5 | 8.00 | C5H11NO2S |
| Phenylalanine | 165.2 | 9 | 4.44 | C9H11NO2 |
| D-Mannose | 180.156 | 6 | 6.67 | C6H12O6 |
| D-Glucose | 180.156 | 6 | 6.67 | C6H12O6 |
| D-Galactose | 180.156 | 6 | 6.67 | C6H12O6 |
| L-Fucose | 164.16 | 6 | 6.67 | C6H12O5 |
| D-Fructose | 180.156 | 6 | 6.67 | C6H12O6 |
| D-Glucuronic Acid | 194.139 | 6 | 6.67 | C6H10O7 |
| Glycerol | 92.094 | 3 | 13.33 | C3H8O3 |
| m-Inositol | 180.16 | 6 | 6.67 | C6H12O6 |
| Mannitol | 182.172 | 6 | 6.67 | C6H14O6 |
| N-Acetyl-D-Glucosamine | 221.21 | 8 | 5.00 | C8H15NO6 |
| Taurine | 125.14 | 2 | 20.00 | C2H7NO3S |
| Maleic Acid | 116.1 | 4 | 10.00 | C4H4O4 |
| Xylitol | 152.15 | 5 | 8.00 | C5H12O5 |
| Acetaldehyde | 44.05 | 2 | 20.00 | C2H4O |
| Sodium Butyrate | 110 | 4 | 10.00 | C4H7NaO2 |
| Maltitol | 344.3 | 12 | 3.33 | C12H24O11 |
| Sodium Gluconate | 218 | 6 | 6.67 | C6H11NaO7 |
| Urea | 60 | 1 | 40.00 | CH4N2O |
| Sorbitol | 182 | 6 | 6.67 | C6H14O6 |
| Oxalacetic Acid | 132 | 4 | 10.00 | C4H4O5 |
| Dulcitol | 182.172 | 6 | 6.67 | C6H14O6 |
| Valeric acid | 102 | 5 | 8.00 | C5H10O2 |
| L-Lyxose | 150 | 5 | 8.00 | C5H10O5 |
| Ethylene Glycol | 62 | 2 | 20.00 | C2H6O2 |
| Raffinose | 594 | 18 | 2.22 | C18H32O16 · 5H2O |
| D+Galactosamine | 179.17 | 6 | 6.67 | C6H13NO5 |
| Lactulose | 342 | 12 | 3.33 | C12H22O11 |
| D+Cellobiose | 342 | 12 | 3.33 | C12H22O11 |
| Glyoxylic Acid | 74 | 2 | 20.00 | C2H2O3 |
| L-Sorbose | 180 | 6 | 6.67 | C6H12O6 |
| Sodium Glycolate | 76.05 | 2 | 20.00 | C2H4O3 |
| Propanol | 60 | 3 | 13.33 | C3H8O |
| Ethanol | 46 | 2 | 20.00 | C2H6O |
| Methanol | 32 | 1 | 40.00 | CH3OH |
| Valine | 117.148 | 5 | 8.00 | C5H11NO2 |
| Norvaline | 117.148 | 5 | 8.00 | C5H11NO2 |
| Tyrosine | 181.191 | 9 | 4.44 | C9H11NO3 |
| Cystine | 249.29 | 6 | 6.67 | C6H12N2O4S2 |
| N-Acetyl-D-Galactosamine | 221.21 | 8 | 5.00 | C8H15NO6 |
| Sodium Formate | 68.007 | 1 | 40.00 | HCOONa |
| Beta-Alanine | 89.09 | 3 | 13.33 | C3H7NO2 |
| 4-Aminobenzoic Acid | 137.138 | 7 | 5.71 | C7H7NO2 |
| Adenine | 135.13 | 5 | 8.00 | C5H5N5 |

**Table S2: Osmolyte Measurements Under Different Growth Conditions**

| | GR [/h] | OD600 | Glu [nmol/OD600/mL] | Gln [nmol/OD600/mL] | DW [mg/OD600] | Glu [% DW] | Gln [% DW] |
|---|---|---|---|---|---|---|---|
| GlcNAc | 0.86 | 0.25 | 169 | 25 | 0.52 | 4.8 | 0.71 |
| 10 mM | 0.86 | 0.37 | 172 | 26 | 0.52 | 4.9 | 0.73 |
| | 0.86 | 0.49 | 165 | 26 | 0.52 | 4.7 | 0.74 |
| Glycerol | 0.40 | 0.26 | 222 | 9 | 0.58 | 5.6 | 0.22 |
| 20 mM | 0.40 | 0.37 | 229 | 8 | 0.58 | 5.8 | 0.20 |
| | 0.40 | 0.49 | 236 | 7 | 0.58 | 6.0 | 0.17 |
| Galactose | 0.29 | 0.31 | 194 | 3 | 0.59 | 4.8 | 0.06 |
| 10 mM | 0.29 | 0.39 | 205 | 2 | 0.59 | 5.1 | 0.04 |
| | 0.29 | 0.50 | 215 | 0 | 0.59 | 5.3 | 0.01 |
| Glucose | 0.83 | 0.27 | 175 | 23 | 0.52 | 5.0 | 0.64 |
| 10 mM | 0.83 | 0.38 | 172 | 22 | 0.52 | 4.9 | 0.63 |
| | 0.83 | 0.50 | 185 | 21 | 0.52 | 5.2 | 0.59 |

**Table S3: *Vibrio splendidus* sp. 1A01 Biomass Reaction**

| Molecular Type | Macromolecular Percentage | Metabolite (BioCyc) | $mol_{mono}/mol_{macro}$ | Formula | C | H | N | O | P | S | MW (g/mol) | Molecular Correction | $MW_{corrected}$ (g/mol) | $g_{mono}/mol_{macro}$ | $g_{macro}/mol_{macro}$ | $g_{mono}/g_{macro}$ | $g_{mono}/g_{CDW}$ | $mmol_{mono}/g_{CDW}$ | Coefficient in Model | Metabolite Name in Model |
|---|---|---|---|---|---|---|---|---|---|---|---|---|---|---|---|---|---|---|---|---|
| **Reactants** | | | | | | | | | | | | | | | | | | | | |
| Protein | 0.58 | L-ALPHA-ALANINE | 0.083 | C3H7NO2 | 3 | 7 | 1 | 2 | 0 | 0 | 89.094 | H2O | 71.079 | 5.89039768 | 110.9483849 | 0.053091333 | 3.08% | 0.433 | -0.433222 | L-ALPHA-ALANINE[c] |
| | | ARG | 0.042 | C6H15N4O2 | 6 | 15 | 4 | 2 | 0 | 0 | 175.212 | H2O | 157.197 | 6.581298862 | | 0.059318564 | 3.44% | 0.219 | -0.218864 | ARG[c] |
| | | ASN | 0.044 | C4H8N2O3 | 4 | 8 | 2 | 3 | 0 | 0 | 132.119 | H2O | 114.104 | 5.022755035 | | 0.045271096 | 2.63% | 0.230 | -0.230117 | ASN[c] |
| | | L-ASPARTATE | 0.057 | C4H6NO4 | 4 | 6 | 1 | 4 | 0 | 0 | 132.095 | H2O | 114.08 | 6.478906397 | | 0.05839568 | 3.39% | 0.297 | -0.296892 | L-ASPARTATE[c] |
| | | CYS | 0.010 | C3H7NO2S | 3 | 7 | 1 | 2 | 0 | 1 | 121.16 | H2O | 103.145 | 1.000625945 | | 0.009018842 | 0.52% | 0.051 | -0.050714 | CYS[c] |
| | | GLN | 0.045 | C5H10N2O3 | 5 | 10 | 2 | 3 | 0 | 0 | 146.146 | H2O | 128.131 | 5.768962313 | | 0.051996812 | 3.02% | 0.235 | | |
| | | GLT | 0.065 | C5H8NO4 | 5 | 8 | 1 | 4 | 0 | 0 | 146.122 | H2O | 128.107 | 8.354022437 | | 0.075296476 | 4.37% | 0.341 | | |
| | | GLY | 0.067 | C2H5NO2 | 2 | 5 | 1 | 2 | 0 | 0 | 75.067 | H2O | 57.052 | 3.849177042 | | 0.034693403 | 2.01% | 0.353 | -0.352699 | GLY[c] |
| | | HIS | 0.021 | C6H9N3O2 | 6 | 9 | 3 | 2 | 0 | 0 | 155.157 | H2O | 137.142 | 2.929982623 | | 0.02640852 | 1.53% | 0.112 | -0.111687 | HIS[c] |
| | | ILE | 0.063 | C6H13NO2 | 6 | 13 | 1 | 2 | 0 | 0 | 131.175 | H2O | 113.16 | 7.17634905 | | 0.06468187 | 3.75% | 0.332 | -0.331526 | ILE[c] |
| | | LEU | 0.101 | C6H13NO2 | 6 | 13 | 1 | 2 | 0 | 0 | 131.175 | H2O | 113.16 | 11.46791712 | | 0.103362632 | 6.00% | 0.530 | -0.529784 | LEU[c] |
| | | LYS | 0.054 | C6H15N2O2 | 6 | 15 | 2 | 2 | 0 | 0 | 147.198 | H2O | 129.183 | 6.930437572 | | 0.062465421 | 3.62% | 0.280 | -0.280454 | LYS[c] |
| | | MET | 0.027 | C5H11NO2S | 5 | 11 | 1 | 2 | 0 | 1 | 149.214 | H2O | 131.199 | 3.489488074 | | 0.031451454 | 1.82% | 0.139 | -0.139040 | MET[c] |
| | | PHE | 0.042 | C9H11NO2 | 9 | 11 | 1 | 2 | 0 | 0 | 165.192 | H2O | 147.177 | 6.182456903 | | 0.055723722 | 3.23% | 0.220 | -0.219598 | PHE[c] |
| | | PRO | 0.038 | C5H9NO2 | 5 | 9 | 1 | 2 | 0 | 0 | 115.132 | H2O | 97.117 | 3.682243888 | | 0.033188801 | 1.92% | 0.198 | -0.198209 | PRO[c] |
| | | SER | 0.071 | C3H7NO3 | 3 | 7 | 1 | 3 | 0 | 0 | 105.093 | H2O | 87.078 | 6.151030842 | | 0.055440472 | 3.22% | 0.369 | -0.369272 | SER[c] |
| | | THR | 0.056 | C4H9NO3 | 4 | 9 | 1 | 3 | 0 | 0 | 119.12 | H2O | 101.105 | 5.655927755 | | 0.050978009 | 2.96% | 0.292 | -0.292441 | THR[c] |
| | | TRP | 0.012 | C11H12N2O2 | 11 | 12 | 2 | 2 | 0 | 0 | 204.229 | H2O | 186.214 | 2.273019347 | | 0.020487178 | 1.19% | 0.064 | -0.063811 | TRP[c] |
| | | TYR | 0.031 | C9H11NO3 | 9 | 11 | 1 | 3 | 0 | 0 | 181.191 | H2O | 163.176 | 4.982378949 | | 0.044907179 | 2.60% | 0.160 | -0.159620 | TYR[c] |
| | | VAL | 0.071 | C5H11NO2 | 5 | 11 | 1 | 2 | 0 | 0 | 117.148 | H2O | 99.133 | 7.081007048 | | 0.063822534 | 3.70% | 0.373 | -0.373408 | VAL[c] |
| DNA | 0.027265982 | DATP | 0.280 | C10H12N5O12P3 | 10 | 12 | 5 | 12 | 3 | 0 | 487.151 | Ppi | 312.202 | 87.37512001 | 307.4682035 | 0.284176116 | 0.77% | 0.0248 | -0.024818 | DATP[c] |
| | | DCTP | 0.220 | C9H12N3O13P3 | 9 | 10 | 3 | 13 | 3 | 0 | 461.109 | Ppi | 286.16 | 62.99318332 | | 0.204877066 | 0.56% | 0.0195 | -0.019521 | DCTP[c] |
| | | DGTP | 0.220 | C10H12N5O13P3 | 10 | 12 | 5 | 13 | 3 | 0 | 503.15 | Ppi | 328.201 | 72.24778361 | | 0.234976439 | 0.64% | 0.0195 | -0.019521 | DGTP[c] |
| | | TTP | 0.280 | C10H13N2O14P3 | 10 | 13 | 2 | 14 | 3 | 0 | 478.136 | Ppi | 303.187 | 84.85211661 | | 0.275970379 | 0.75% | 0.0248 | -0.024818 | TTP[c] |
| RNA | 0.193 | CTP | 0.213 | C9H12N3O14P3 | 9 | 12 | 3 | 14 | 3 | 0 | 479.124 | Ppi | 304.175 | 64.66664902 | 322.91633 | 0.200258219 | 3.86% | 0.1271 | -0.127064 | CTP[c] |
| | | GTP | 0.302 | C10H12N5O14P3 | 10 | 12 | 5 | 14 | 3 | 0 | 519.149 | Ppi | 344.2 | 103.7971504 | | 0.321436672 | 6.20% | 0.1802 | -0.180236 | GTP[c] |
| | | UTP | 0.217 | C9H11N2O15P3 | 9 | 11 | 2 | 15 | 3 | 0 | 480.108 | Ppi | 305.159 | 66.23766802 | | 0.205123315 | 3.96% | 0.1297 | -0.129732 | UTP[c] |
| | | ATP | 0.269 | C10H12N5O13P3 | 10 | 12 | 5 | 13 | 3 | 0 | 503.15 | Ppi | 328.201 | 88.21486253 | | 0.273181795 | 5.27% | 0.1606 | | |
| Murein | 0.021929521 | CPD0-2278 | 1 | C77H117N15O40 | 77 | 117 | 15 | 40 | 0 | 0 | 1892.848 | | 1892.848 | 1892.848 | 1892.848 | 1 | 2.19% | 0.012 | -0.011585 | CPD0-2278[p] |
| LPS | 0.029850831 | KDO2-LIPID-IVA | 1 | C84H148N2O37P2 | 84 | 148 | 2 | 37 | 2 | 0 | 1840.033 | | 1840.033 | 1840.033 | 1840.033 | 1 | 2.99% | 0.016222987 | -0.016223 | KDO2-LIPID-IVA[p] |
| Lipids | 0.079963539 | CPD-12819 | 0.4590 | C37H74N1O8P1 | 37 | 74 | 1 | 8 | 1 | 0 | 691.972 | | 691.972 | 317.615148 | 689.790688 | 0.460451487 | 3.68% | 0.053 | -0.018598 | CPD-12819[c] |
| | | | | | | | | | | | | | | | | | | | -0.034611 | CPD-12819[p] |
| | | CPD-17086 | 0.5410 | C37H70N1O8P1 | 37 | 70 | 1 | 8 | 1 | 0 | 687.94 | | 687.94 | 372.17554 | | 0.539548513 | 4.31% | 0.063 | -0.021934 | CPD-17086[c] |
| | | | | | | | | | | | | | | | | | | | -0.040781 | CPD-17086[p] |
| Inorganic ions | 0.008755132 | K+ | 0.7143 | K | | | | | | | 38.9637 | | 38.9637 | 27.83121429 | 42.21914254 | 0.659208421 | 0.58% | 0.148 | -0.148124 | K+[c] |
| | | AMMONIUM | 0.0476 | H4N | 0 | 4 | 1 | 0 | 0 | 0 | 18.039 | | 18.039 | 0.859 | | 0.020346221 | 0.02% | 0.010 | -0.009875 | AMMONIUM[c] |
| | | MG+2 | 0.0317 | Mg | | | | | | | 23.985 | | 23.985 | 0.761428571 | | 0.01803515 | 0.02% | 0.007 | -0.006583 | MG+2[c] |
| | | CA+2 | 0.0190 | Ca | | | | | | | 39.9626 | | 39.9626 | 0.761192381 | | 0.018029556 | 0.02% | 0.004 | -0.003950 | CA+2[c] |
| | | FE+2 | 0.0286 | Fe | | | | | | | 55.9349 | | 55.9349 | 1.59814 | | 0.037853445 | 0.03% | 0.006 | -0.005925 | FE+2[c] |
| | | FE+3 | 0.0286 | Fe | | | | | | | 55.9349 | | 55.9349 | 1.59814 | | 0.037853445 | 0.03% | 0.006 | -0.005925 | FE+3[c] |
| | | CU+2 | 0.0127 | Cu | | | | | | | 63.546 | | 63.546 | 0.806933333 | | 0.019112973 | 0.02% | 0.003 | -0.002633 | CU+2[c] |
| | | MN+2 | 0.0127 | Mn | | | | | | | 54.938 | | 54.938 | 0.697625397 | | 0.016523912 | 0.01% | 0.003 | -0.002633 | MN+2[c] |
| | | CPD-3 | 0.0127 | MoO4 | 0 | 0 | 0 | 4 | 0 | 0 | 159.94 | | 159.94 | 2.030984127 | | 0.048105764 | 0.04% | 0.003 | -0.002633 | CPD-3[c] |
| | | CO+2 | 0.0127 | Co | | | | | | | 58.9332 | | 58.9332 | 0.748358095 | | 0.017725564 | 0.02% | 0.003 | -0.002633 | CO+2[c] |
| | | ZN+2 | 0.0127 | Zn | | | | | | | 63.9291 | | 63.9291 | 0.811798095 | | 0.019228199 | 0.02% | 0.003 | -0.002633 | ZN+2[c] |
| | | CL- | 0.0190 | Cl | | | | | | | 34.9689 | | 34.9689 | 0.666074286 | | 0.015776594 | 0.01% | 0.004 | -0.003950 | CL-[c] |
| | | SULFATE | 0.0159 | O4S | 0 | 0 | 0 | 4 | 0 | 1 | 96.062 | | 96.062 | 1.524793651 | | 0.036116168 | 0.03% | 0.003 | -0.003292 | SULFATE[c] |
| | | Pi | 0.0159 | HO4P | 0 | 1 | 0 | 4 | 1 | 0 | 95.978 | | 95.978 | 1.523460317 | | 0.036084587 | 0.03% | 0.003 | | |
| Soluble pools | 0.002834995 | CO-A | 0.000576 | C21H32N7O16P3S | 21 | 32 | 7 | 16 | 3 | 1 | 763.508 | | 763.508 | | | | | 0.000480 | -0.000480 | CO-A[c] |
| | | NAD | 0.001831 | C21H26N7O14P2 | 21 | 26 | 7 | 14 | 2 | 0 | 662.422 | | 662.422 | | | | | 0.001527 | -0.001527 | NAD[c] |
| | | NADP | 0.000447 | C21H25N7O17P3 | 21 | 25 | 7 | 17 | 3 | 0 | 740.385 | | 740.385 | | | | | 0.000372 | -0.000372 | NADP[c] |
| | | FAD | 0.000223 | C27H31N9O15P2 | 27 | 31 | 9 | 15 | 2 | 0 | 783.541 | | 783.541 | | | | | 0.000186 | -0.000186 | FAD[c] |
| | | THF-GLU-N | 0.000223 | C19H21N7O6 | 19 | 21 | 7 | 6 | 0 | 0 | 443.42 | | 443.42 | | | | | 0.000186 | -0.000186 | THF-GLU-N[c] |
| | | METHYLENE-THF-GLU-N | 0.000223 | C20H21N7O6 | 20 | 21 | 7 | 6 | 0 | 0 | 455.431 | | 455.431 | | | | | 0.000186 | -0.000186 | METHYLENE-THF-GLU-N[c] |
| | | 5-METHYL-THF-GLU-N | 0.000223 | C20H24N7O6 | 20 | 24 | 7 | 6 | 0 | 0 | 458.455 | | 458.455 | | | | | 0.000186 | -0.000186 | 5-METHYL-THF-GLU-N[c] |
| | | 10-FORMYL-DIHYDROFOLATE-GLU-N | 0.000223 | C20H21N7O7 | 20 | 21 | 7 | 7 | 0 | 0 | 471.43 | | 471.43 | | | | | 0.000186 | -0.000186 | 10-FORMYL-DIHYDROFOLATE-GLU-N[c] |
| | | THIAMINE-PYROPHOSPHATE | 0.000223 | C12H16N4O7P2S | 12 | 16 | 4 | 7 | 2 | 1 | 422.295 | | 422.295 | | | | | 0.000186 | -0.000186 | THIAMINE-PYROPHOSPHATE[c] |
| | | CPD-9956 | 0.000223 | C49H76O4 | 49 | 76 | 0 | 4 | 0 | 0 | 729.143 | | 729.143 | | | | | 0.000186 | -0.000186 | CPD-9956[c] |
| | | PYRIDOXAL_PHOSPHATE | 0.000223 | C8H8NO6P | 8 | 8 | 1 | 6 | 1 | 0 | 245.127 | | 245.127 | | | | | 0.000186 | -0.000186 | PYRIDOXAL_PHOSPHATE[c] |
| | | PROTOHEME | 0.000223 | C34H30FeN4O4 | 34 | 30 | 4 | 4 | 0 | 0 | 614.5729 | | 614.5729 | | | | | 0.000186 | -0.000186 | PROTOHEME[c] |
| | | ENTEROBACTIN | 0.000223 | C30H27N3O15 | 30 | 27 | 3 | 15 | 0 | 0 | 669.552 | | 669.552 | | | | | 0.000186 | -0.000186 | ENTEROBACTIN[c] |
| | | UNDECAPRENYL-DIPHOSPHATE | 0.000055 | C55H89O7P2 | 55 | 89 | 0 | 7 | 2 | 0 | 924.258 | | 924.258 | | | | | 0.000046 | -0.000046 | UNDECAPRENYL-DIPHOSPHATE[c] |
| | | CHORISMATE | 0.000223 | C10H8O6 | 10 | 8 | 0 | 6 | 0 | 0 | 224.168 | | 224.168 | | | | | 0.000186 | -0.000186 | CHORISMATE[c] |
| | | S-ADENOSYLMETHIONINE | 0.000223 | C15H23N6O5S | 15 | 23 | 6 | 5 | 0 | 1 | 399.452 | | 399.452 | | | | | 0.000186 | -0.000186 | S-ADENOSYLMETHIONINE[c] |
| | | RIBOFLAVIN | 0.000223 | C17H20N4O6 | 17 | 20 | 4 | 6 | 0 | 0 | 376.369 | | 376.369 | | | | | 0.000186 | -0.000186 | RIBOFLAVIN[c] |
| Osmolytes | 0.0564 | GLN | 0.109929078 | C5H10N2O3 | 5 | 10 | 2 | 3 | 0 | 0 | 146.146 | | 146.146 | 16.06569504 | 146.1246383 | 0.109945148 | 0.62% | 0.042 | | |
| | | GLT | 0.890070922 | C5H8NO4 | 5 | 8 | 1 | 4 | 0 | 0 | 146.122 | | 146.122 | 130.0589433 | | 0.890054852 | 5.02% | 0.344 | | |
| RNA + GAM | | | | | | | | | | | | | | | | | | 17.470646 | -17.470646 | ATP[c] |
| Protein + osmolytes | | | | | | | | | | | | | | | | | | 0.277799 | -0.277799 | GLN[c] |
| Protein + osmolytes | | | | | | | | | | | | | | | | | | 0.684445 | -0.684445 | GLT[c] |
| GAM - protein polymerization | | WATER | | H2O | 0 | 2 | 0 | 1 | 0 | 0 | 18.015 | | | | | | | 12.082370 | -12.082370 | WATER[c] |
| **Products** | | | | | | | | | | | | | | | | | | | | |
| GAM | | | | | | | | | | | | | | | | | | 17.310000 | 17.310000 | ADP[c] |
| GAM | | | | | | | | | | | | | | | | | | 17.310000 | 17.310000 | PROTON[c] |
| Sum of dNTPs | | | | | | | | | | | | | | | | | | 0.0887 | | |
| Sum of NTPs | | | | | | | | | | | | | | | | | | 0.5977 | | |
| Sum of amino acids | | | | | | | | | | | | | | | | | | 5.228 | | |
| NTPs + dNTPs | | PPI | | HO7P2 | 0 | 1 | 0 | 7 | 2 | 0 | 174.949 | | | | | | | 0.686357 | 0.686357 | PPI[c] |
| GAM - inorganic ions | | | | | | | | | | | | | | | | | | 17.306708 | 17.306708 | Pi[c] |

Adapted from Sup. Info. 3 in Feist, A. M. *et al.* A genome-scale metabolic reconstruction for Escherichia coli K-12 MG1655 that accounts for 1260 ORFs and thermodynamic information. Mol. Syst. Biol. (2007)

**Table S4: Calculation of Uptake and Secretion Fluxes**

| Carbon source | Number of carbon atoms | Growth rate ($hr^{-1}$) | 1/Yield (mM substrate/OD) | Acetate excretion yield (mM/OD) | Carbon yield (mM C/OD) | Carbon utilization rate (mM C/OD/h) | Carbon utilization rate (mmol C/gCDW/h) | OD-to-CDW conversion (mgCDW/OD *mL) | Acetate excretion rate (mmol/gCDW/h) | Carbon uptake rate (mmol/gCDW/h) |
|---|---|---|---|---|---|---|---|---|---|---|
| GlcNAc | 8 | 0.82 | 5.8 | 8.6 | 29.2 | 23.94 | 45.93 | 0.52 | 13.53 | 9.12 |
| Glucose | 6 | 0.79 | 5.5 | 2.2 | 28.6 | 22.59 | 43.01 | 0.53 | 3.31 | 8.27 |
| Galactose | 6 | 0.29 | 6.3 | 0 | 37.8 | 10.96 | 18.49 | 0.59 | 0.00 | 3.08 |
| Pyruvate | 3 | 0.22 | 22.1 | 3.8 | 58.7 | 12.91 | 21.44 | 0.60 | 1.39 | 8.07 |

### Document S1: Minimal MBL Medium

**4x Seawater**

This is the base for all defined media.

| Component | Amount (per L) | FW (g/mol) | Concentration (mM) |
|---|---|---|---|
| NaCl | 80 g | 58.44 | 1369 |
| $MgCl_2$-$6H_2O$ | 12 g | 203.20 | 59 |
| $CaCl_2$-$2H_2O$ | 0.60 g | 147.02 | 4 |
| KCl | 2.0 g | 74.56 | 27 |

Make 1 L and filter sterilize through 0.2 µm.

**1000x Trace Minerals (per L)**

Add to recapitulate the ionic composition of seawater.

Dissolve in 20 mM HCl (to avoid precipitate):

| Substance | mg / L |
|---|---|
| $FeSO_4$ * $7H_2O$ | 2100 |
| $H_3BO_3$ | 30 |
| $MnCl_2$ * $4H_2O$ | 100 |
| $CoCl_2$ * $6H_2O$ | 190 |
| $NiCl_2$ * $6H_2O$ | 24 |
| $CuCl_2$ * $2H_2O$ | 2 |
| $ZnSO_4$ * $7H_2O$ | 144 |
| $Na_2MoO_4$ * $2H_2O$ | 36 |
| $NaVO_3$ | 25 |
| $NaWO_4$ $2H_2O$ | 25 |
| $Na_2SeO_3$ $5H_2O$ | 6 |

**Note*: $NaVO_3$ and $NaWO_4$ * $2H_2O$ should only be opened in the hood.

Filter sterilize through 0.2 µm.
Store at 4 C in the dark. Stable for a couple months.
For long-term storage, freeze at -20 C in aliquots of 1 mL.

**1000x Vitamins**

Dissolve in 10 mM MOPS, pH 7.2:

| Substance | mg/L |
|---|---|
| Riboflavin | 100 |
| D-Biotin | 30 |
| Thiamine hydrochloride | 100 |
| L-ascorbic acid | 100 |
| Ca-d- pantothenate | 100 |
| Folate | 100 |
| Nicotinate | 100 |
| 4-aminobenzoic acid | 100 |
| pyridoxine HCl | 100 |
| Lipoic acid | 100 |

| | |
|---|---|
| NAD | 100 |
| Thiamin pyrophosphate | 100 |
| Cyanocobalamin | 10 |

Titrate with a couple of drops of 5 M NaOH to avoid precipitate.
Filter sterilize through 0.2 μm.
Store at 4 C in the dark. Stable for a couple months.
For long-term storage, freeze at -20 C in 1 mL aliquots.

**Nitrogen source**
**1 M ammonium chloride (100x)**

Dissolve 2.14 g $NH_4Cl$ in 40 mL distilled water.
Filter sterilize through 0.2 μm.

**Phosphorus source**
**0.5 M phosphate dibasic (500x)**

Dissolve 2.84 g $Na_2HPO_4$ in 40 mL distilled water.
Filter sterilize through 0.2 μm.

**Sulfur source**
**1 M sodium sulfate (1000x)**

Dissolve 5.68 g $Na_2SO_4$ in 40 mL distilled water.
Filter sterilize through 0.2 μm.

**HEPES buffer:**
**1 M HEPES buffer (20x), pH 8.2**

Per L
260.29 g HEPES sodium salt
Dissolve in 750 mL distilled water. Adjust pH to 8.2 with concentrated HCl and constant stirring.
Bring final volume to 1 L with water. Filter sterilize through 0.2 μm. Store at 4 C.

**Basic recipe for medium (40 mL)**

0.04 mL vitamins
0.04 mL trace metals
0.04 mL 1000x sodium sulfate
0.08 mL of 500x phosphate dibasic
0.4 mL of 100x ammonium chloride

2 mL of 20x GlcNAc
2 mL of 20x HEPES buffer, pH 8.2
10 mL of 4x seawater
25.4 mL of autoclaved $ddH_2O$
